# Recurrence of cancer cell states across diverse tumors and their interactions with the microenvironment

**DOI:** 10.1101/2021.12.20.473565

**Authors:** Dalia Barkley, Reuben Moncada, Maayan Pour, Deborah A. Liberman, Ian Dryg, Gregor Werba, Wei Wang, Maayan Baron, Anjali Rao, Bo Xia, Gustavo S. França, Alejandro Weil, Deborah F. Delair, Cristina Hajdu, Amanda W. Lund, Iman Osman, Itai Yanai

## Abstract

While genetic tumor heterogeneity has long been recognized, recent work has revealed significant variation among cancer cells at the epigenetic and transcriptional levels. Profiling tumors at the single-cell level in individual cancer types has shown that transcriptional heterogeneity is organized into cancer cell states, implying that diverse cell states may represent stable and functional units with complementary roles in tumor maintenance and progression. However, it remains unclear to what extent these states span tumor types, constituting general features of cancer. Furthermore, the role of cancer cell states in tumor progression and their specific interactions with cells of the tumor microenvironment remain to be elucidated. Here, we perform a pan-cancer single-cell RNA-Seq analysis across 15 cancer types and identify a catalog of 16 gene modules whose expression defines recurrent cancer cell states, including ‘stress’, ‘interferon response’, ‘epithelial-mesenchymal transition’, ‘metal response’, ‘basal’ and ‘ciliated’. Using mouse models, we find that induction of the interferon response module varies by tumor location and is diminished upon elimination of lymphocytes. Moreover, spatial transcriptomic analysis further links the interferon response in cancer cells to T cells and macrophages in the tumor microenvironment. Our work provides a framework for studying how cancer cell states interact with the tumor microenvironment to form organized systems capable of immune evasion, drug resistance, and metastasis.

## Introduction

Transcriptional heterogeneity in cancer is increasingly recognized as a driver of tumor progression, metastasis and treatment failure^1–5^. Single-cell RNA-Sequencing (scRNA-Seq) has enabled the unbiased transcriptomic profiling of individual tumor cells and has revealed a striking amount of heterogeneity among malignant cells of the same tumor^6–12^. Furthermore, evidence has emerged suggesting that transcriptional heterogeneity is organized into modules of co-expressed genes^13^. Data from glioblastoma^6,7^, oligodendroglioma^14^, astrocytoma^15^, head and neck cancer^10^ and melanoma^16,17^ among others, indicates that, within a tumor, cancer cells are heterogeneous in their degree of differentiation, ranging from stem- or progenitor-like to fully differentiated. These studies performed in a variety of cancer types have also shown the existence of cancer cell states related to stress response, interferon response, and hypoxia^6–10^. While certain states have been found in multiple studies, a general catalog of cell states across cancer types remains to be established. Such a coherent framework - if it exists - would allow us to search for common themes across cancer types and to understand how tumors are organized independently of their origin.

Beyond malignant cells, tumors are composed of a complex microenvironment including immune and stromal cells, which also play critical roles in tumorigenesis^18^. In particular, the clinical success of immunotherapy across multiple cancers^19–22^ hints at commonalities in the interactions between cancer cells and the tumor microenvironment (TME). Causative links have been drawn between specific elements of the TME and cancer cell states^10,23,24^. In one study of head and neck cancer, a population of partial epithelial-mesenchymal transition (pEMT) cancer cells at the leading edge of tumors was shown to interact with cancer-associated fibroblasts and mediate invasion^10^. In glioblastoma and triple-negative breast cancer, factors of the TME appear to induce malignant cells to adopt a stem-like state^23,24^. These works point to a need for a systematic analysis of cancer cell states, with a particular focus on the relation to the non-malignant cell types of the TME.

Here, we characterize recurrent cancer cell states and their relationship with the TME by systematically assaying 15 cancer types to identify a catalog of recurrent cancer cell states using a gene-centric approach. Analyzing scRNA-Seq data from previously published data as well as newly collected tumors, we identified 16 coherent gene modules and quantified their expression in malignant cells of each sample. This catalog includes modules present in all studied tumors, as well as others that are specific to particular sets of cancer types. To further study the cancer cells states, we used experimental models to perturb the tumor microenvironment and test for differential effects on the cancer cell states. While some of the states are related to known aspects of cancer biology, we present evidence that these processes are heterogeneously deployed by cells of the same tumor, and that this heterogeneity recurs across a wide range of cancer types. A detailed analysis of the interferon response module further led us to study its dependencies *in vivo* in the context of TME perturbations and to establish its proximity to macrophages and T cells across cancer types. Overall, the catalog of cancer cell states is a coherent representation of the makeup of a tumor, and provides a framework for the analysis and testing of the features of tumorigenesis.

### Recurring gene modules across diverse cancer types

We collected 19 fresh primary untreated patient tumors spanning 9 cancer types immediately after surgery (Supplementary Table 1). We dissociated each tumor to obtain a single-cell suspension, and processed for scRNA-Seq without prior sorting to ensure an unbiased assessment of the tumor cellular composition. Our tumor collection included 9 cancer types: carcinoma of the ovary (OVCA), endometrium (UCEC), breast (BRCA), prostate (PRAD), kidney (KIRC), liver (LIHC), colon (COAD) and pancreas (PDAC), as well as one non-epithelial cancer type, gastrointestinal stromal tumor (GIST) (Fig. 1a). We first identified the malignant cells in our dataset by analyzing the transcriptomes using a combination of marker genes, singleR annotation^25^, and inferred copy number variation^26^, and controlling for the possibility of doublets (Fig. 1b-c, Extended Data Fig. 1-2, see Methods). Across our samples, we annotated 9,036 malignant and 18,546 non-malignant cells (Supplementary Table 2). To extend our dataset, we also performed this analysis in tumors from prior publications, including additional PDAC^27^ and LIHC^28^, as well as additional tumor types: cholangiocarcinoma (CHCA)^29^, lung adenocarcinoma (LUAD)^30^, head and neck squamous cell carcinoma (HNSC)^10^, skin squamous cell carcinoma (SKSC)^31^, glioblastoma multiforme (GBM)^7^ and oligodendroglioma (OGD)^14^, resulting in a total of 19,942 malignant cells from 62 untreated primary tumors spanning 15 cancer types (Fig. 1a).

**Figure 1:**
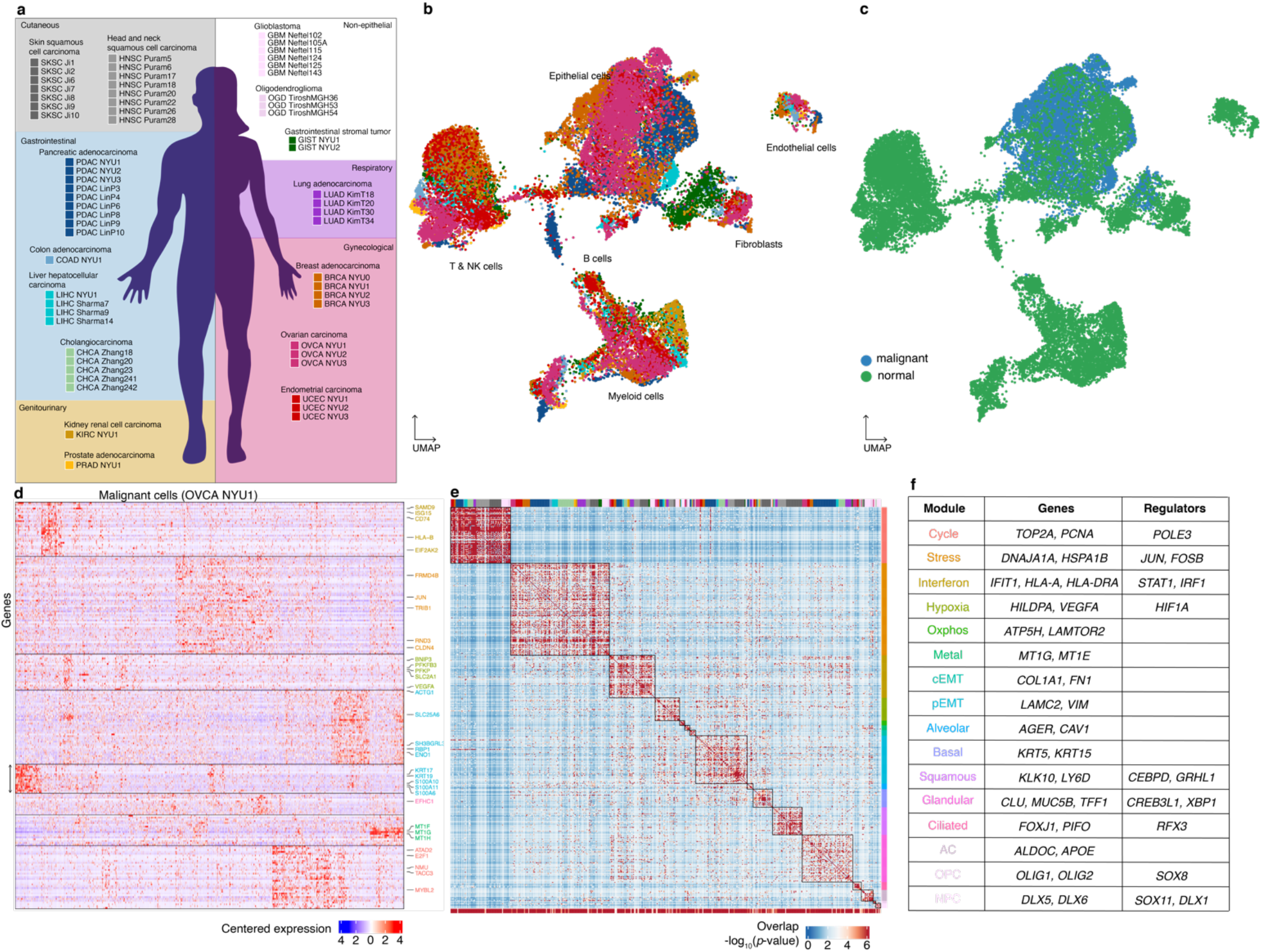
A catalog of recurrent cancer gene modules. **a**. Tumors collected in this study and tumors included from previous reports for joint analysis. The background color indicates the organ system of origin: cutaneous (grey), gastrointestinal (blue), gynecological (red) and genitourinary (yellow). White background indicates non-epithelial tumors. **b**. UMAP embedding of cells from 19 tumors collected spanning a total of 9 cancer types, colored as in **a**. **c**. As in **b**., colored by annotation as malignant or non-malignant **d**. Heatmap of expression levels for 241 genes in the malignant cells of the ovarian tumor OVCA NYU1. Genes are ordered by their module membership (horizontal lines) and the colors of the indicated genes correspond to their consensus module annotation described in **e**. **e**. Heatmap of the significance of the overlap between individual tumor modules (hypergeometric test). The bottom bar indicates the significance of the overlap with consensus modules (hypergeometric test). The top bar indicates the identity of the tumor samples, colored as in a. **Extended Data Figure 3h** indicates the significance of the overlap of each consensus module with each tumor specific gene module. **f**. Table of consensus modules, selected genes and putative regulators identified using SCENIC (regulators identified in at least 2 tumors are shown), colored as in **d**. See also **Extended Data Figure 3h** and Supplementary Table 4.

Our approach to defining cell states involved first cataloging the underlying gene modules, following recent work that has identified gene modules as the defining features of cell states^6,7,10,14^. This is a flexible approach since it allows for cells expressing combinations of modules, and thus for the complexity of possible cell states. We analyzed the malignant cells using non-negative matrix factorization (NMF) to identify gene modules as sets of co-expressed genes (Fig. 1d, see Methods). Our method detects groups of genes that are co-expressed within the sample, *i*.*e*. that are expressed coherently in a subset of cells. To search for recurring gene modules across tumors, we then compared the gene composition of the identified modules (Fig. 1e, see Methods). By thus performing the integration at the level of gene modules rather than expression matrices, the impact of technical variation across the samples and studies is limited. Despite the independent identification of these modules in a variety of cancer types – thereby not presuming recurrence – we found that modules obtained in different tumors overlap significantly (Fig. 1e, top bar). Importantly, this finding of recurrence rather than uniqueness would not be the result of batch effects.

The recurrence of the gene modules enabled us to construct a catalog of 16 consensus modules (henceforth ‘modules’; Fig. 1f, Extended Data Fig. 3f, Supplementary Table 3, see Methods), with a median of 37 genes per module. To establish whether a module is significantly present in the population of cancer cells in each tumor, we used the gene set overdispersion metric^32^ (Extended Data Fig. 4a, see Methods). Consistent with Figure 1d, some modules were enriched in specific organ systems (such as the brain or gynecological organs), while others spanned several organ systems and histologies (see below). We also tested for the overdispersion of the modules in independent datasets representing normal epithelia from the fallopian tube^33^, breast^34^ and liver^35^ (Extended Data Fig. 4b) in order to ask whether the modules reflect a reconstitution of the heterogeneity found in normal tissues. For most modules, we found that the overdispersion was lower in normal epithelial samples, suggesting that they are not as differentially expressed in normal tissues, with some exceptions which we detail below. However, the fact that the catalog of modules are indeed detected to some extent in normal tissues suggests that the modules are not specific to cancer, but rather are co-opted from existing ones and expressed more heterogeneously (see Discussion). We also studied the gene composition and cancer type-specificity of the modules, distinguishing modules related to cellular processes (Extended Data Fig. 3a) from those related to cell identity (Extended Data Fig. 3b).

As expected, we recovered a highly recurrent module consisting of cell cycle genes (e.g., *TOP2A, PCNA*), capturing the subset of cancer cells in any tumor that is cycling at the time of sampling. Another process which recurred across tumor types was the stress response (e.g., *JUN, FOS, HSPA1B*), which has been previously described^12,36,37^ and shown to have a role in drug resistance in melanoma^17^. We also present spatial transcriptomics data below (Fig. 5) that provides additional support for the *in vivo* existence of this state among cancer cells in the absence of dissociation.

An interferon response module, which has been identified in metastatic ovarian carcinoma^12^, was detected as widely occuring, showing that interferon response in malignant cells is heterogeneous across a range of solid tumor types. In addition to interferon stimulated genes such as *STAT1* and *IFIT1*, this module contained components of antigen presentation, a well-characterized effect of interferon^38^, with both MHC I genes such as *HLA-A*^*39,40*^ and MHC II genes including *HLA-DRA*. While MHC II expression is classically associated with professional antigen-presenting cells, this pathway has also been shown to be expressed in normal epithelial cells and in cancer cells^41,42^. Interferon response generally functions as a defense response recruiting and activating immune cells, and has been extensively studied in cancer^43– 45^. In this context, interferon ligands may be secreted by cancer cells and dendritic cells (DCs) (for type I interferons, IFNα and IFNβ), or by natural killer (NK) and T cells (for IFNγ). Alternatively, the interferon response may be cancer cell intrinsic, i.e. activated independently from signaling by other cell types; indeed, a recent study comparing gene modules across cancer cell lines *in vitro* also identified an interferon response module^46^, supporting the possibility of a TME-independent response.

Two modules relating to metabolic processes were also found across a range of cancer types: a hypoxia module (e.g., *VEGF, ADM*)^6,47–50^ and an oxidative phosphorylation module (e.g., *ATP5H, LAMTOR2*)^9^. Metabolic adaptation to hypoxia in solid tumors, with increased glycolysis and induction of angiogenesis, has been implicated in cancer progression, drug resistance, invasion and metastasis^51,52^. Nonetheless, recent studies have shown a role for oxidative phosphorylation in several cancer types, suggesting that cancer cells may rely on both glycolysis and oxidative phosphorylation for energy production^53,54^. An additional gene module of metallothionein genes – which we refer to here as a metal-response module – may have a role in proliferation and drug resistance in several cancer types^55–58^.

Another set of modules correspond to cell identity, and appear to be related to the tissue and cell of origin (Extended Data Fig. 3b). The majority of the tumors profiled were of epithelial origin, and accordingly we identified modules overlapping with known epithelial cell type markers: an alveolar module (e.g., *AGER, CAV1*) which was particularly present as expected in LUAD^59–61^, as well as basal (e.g., *KRT5* and *KRT15*), squamous (e.g., *KLK10, LY6D*), and glandular (e.g., *CLU, MUC5B*) cell modules (Extended Data Fig. 3f-g). A module composed of cilium-related genes (e.g., *FOXJ1, PIFO*) was present in gynecological tumors as well as LUAD and GBM. In ovarian and endometrial tumors, this module was present only in the endometrioid samples (OVCA NYU2-3, UCEC NYU2-3), and not in the high grade serous samples (OVCA NYU1, UCEC NYU1), pointing to cilium formation as a characteristic of endometrioid histology. The presence of the module in normal fallopian tube and lung epithelial tissues (Extended Data Fig. 4b-c)^62,63^ suggests that its differential expression in cancer mirrors the heterogeneity of the tissue of origin. The heterogeneous expression of differentiation modules that we observe within tumors may provide a more detailed understanding of tumor architecture from a clinical pathology perspective where each tumor is assessed for grade and histological subtype.

Two of the modules spanning multiple cancer types were related to epithelial-mesenchymal transition (EMT): a complete mesenchymal module (cEMT) (e.g., *COL1A1, FN1*) and a partial mesenchymal module (pEMT) (e.g., *LAMC2, VIM*) lacking canonical mesenchymal markers such as collagen genes^10^. The pEMT module has been recently characterized in HNSC^10^ and SKSC^31^ (Extended Data Fig. 3c-e), but is also found in GBM^7^, suggesting that cells from different lineages converge upon this identity in cancer. We detected the presence of the cEMT module in a minority of samples, but a range of cancer types: mainly PDAC, CHCA, LUAD, and GBM. A recent study has shown that pEMT and cEMT can occur in a range of cancer types^64^, and may represent two pathways converging upon the phenotypic properties conferred by mesenchymal differentiation including migration and drug resistance^11,65^. Using TCGA data^66^, we indeed found that expression of the pEMT gene module is associated with decreased progression-free survival (Extended Data Fig. 16, see Methods)^67,68^.

Finally, we identified three neurological cancer-specific modules that were analogous to those described by Tirosh et al.^14^ and Neftel et al.^7^ (Extended Data Fig. 3c): the astrocyte (AC)-like (e.g., *APOE, ALDOC*), oligodendrocyte progenitor cell (OPC)-like (e.g., *OLIG1, OLIG2*), and neural progenitor cell (NPC)-like (e.g., *DLX1, DLX5*) modules.

The broad incidence of these modules across a range of cancer types highlights redeployment of differentiation programs and distinct expression levels in cancer and normal tissues. Moreover, while many of the genes identified have been implicated in aspects of cancer biology (as discussed above), our single-cell approach enabled us to show that they are heterogeneously expressed among the malignant cells of a tumor (Extended Data Fig. 4a), and generally expressed at higher levels in malignant versus normal epithelial cells (Extended Data Fig. 4d).

To test whether the catalog of 16 modules can also be detected using an independent approach, we used SCENIC^69^, a method that identifies genes that are both correlated in their expression and regulated by the same transcription factor. We found that each module of our catalog had significant overlap with several SCENIC regulons (Extended Data Fig. 3h, Supplementary Table 4, see Methods). For instance, the interferon response module overlapped with several SCENIC regulons annotated with the transcription factors *STAT1* and *IRF1*.

### Defining cancer cell states by gene module expression

Having established the catalog of cancer gene modules, we next sought to understand how they are generally assembled at the level of individual cells. In particular, we asked whether cells are constrained in which modules or combinations of modules they can express. For this, we scored each malignant cell for the expression of each of the modules (Fig. 2a, see Methods). In the SKSC Ji1 sample, for example, expression of basal, squamous and cycling modules was mutually exclusive, but each of these had co-expression with the stress or pEMT module (Fig. 2a, Extended Data Fig. 7a-b). More generally, we found that most cells express a combination of modules, though not all combinations are possible. These results support the notion that, in defining a cancer cell state, it is crucial to examine the complete set of gene modules expressed.

**Figure 2:**
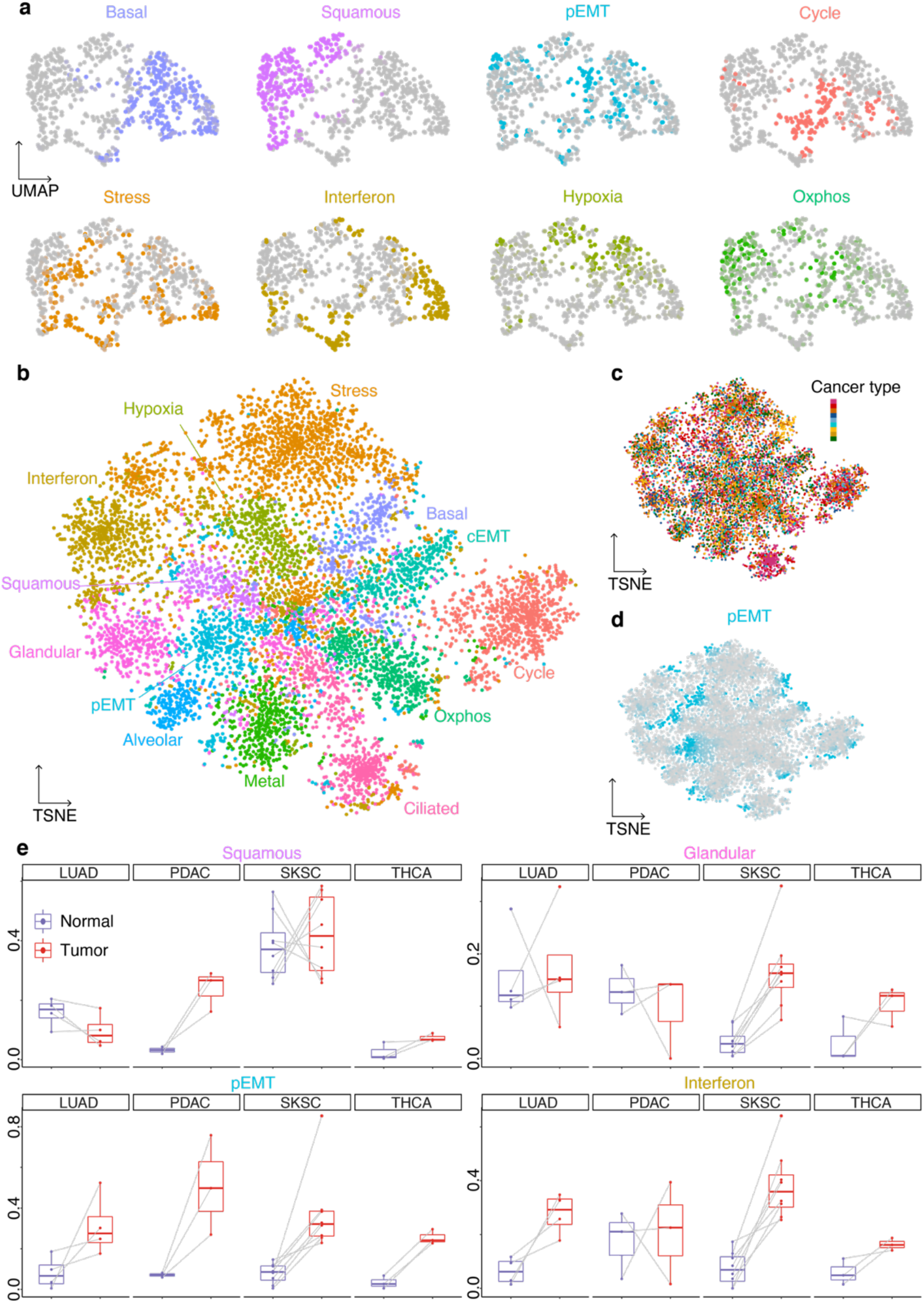
Expression of gene modules underlies cancer cell states. **a**. Gene expression UMAP embedding of malignant cells of SKSC Ji1, colored by module score for the 8 indicated modules. **b**. Module score TSNE embedding of the cancer cells of all 18 tumors, colored by the most high scoring module. **c**. Same as **b**, colored by cancer type, as in **Figure 1a**. **d**. Same as **b**, colored by pEMT module score. **e**. Boxplots of the expression frequency (fraction of cells with module score greater than 0.5) of the squamous, glandular, pEMT and interferon response modules in paired normal and tumor samples.

Since the modules recur across cancer types, we reasoned that the cell module scores could serve as natural axes across which to compare cancer cells of different patients. Figure 2b represents a dimensionality reduction performed on the module scores of cells from 19 different tumors collected at NYU (see Methods). Most notably, the cancer cells in this space do not group by patient or cancer type, but rather by their most highly expressed module (Fig. 2b,c, Extended Data Fig. 6a-c). This is in sharp contrast with the finding that, in gene expression space, cancer cells cluster by patient^36^, and highlights commonalities across cancer types when variation due to individual genes is removed. As described for SKSC Ji1 (Fig. 2a), there is a degree of co-expression between certain modules: for example pEMT is co-expressed with stress and interferon-response (Fig. 2d, Extended Data Fig. 6b). Together with the fact that cells do not form distinct clusters, this supports the view that cancer cell states do not generally represent discrete entities. We did observe, however, discrete clusters corresponding to cells expressing the cycle or cilium module. These clusters are also identified when examining tumors individually in gene expression-based dimensionality reductions, and are therefore not artifacts of the module score dimensionality reduction (Extended Data Fig. 6d).

Since certain modules are also present in non-cancer samples (Extended Data Fig. 4b-c), we asked whether the fraction of cells expressing each module varies between malignant and non-malignant epithelia. For this, we compared the module expression frequency in the malignant cells to those in non-malignant cells from matched samples, both in our dataset and that of Kim et al.^30^, Ji et al.^31^ and Pu et al.^70^ (Fig. 2e and Extended Data Fig. 6f). While for LUAD, SKSC and THCA we compared to the epithelial cells of paired adjacent normal samples^30,31^, in PDAC, non-malignant ductal cells from the same samples served as a paired normal comparison (see Methods).

The pEMT module was expressed at higher frequencies in all three cancer types relative to normal (Fig. 2e), in line with the common occurrence of EMT in epithelial cancers^67^. The interferon response module exhibited increased expression frequency in LUAD and SKSC relative to normal, but was unchanged in PDAC (Fig. 2e). This may be partly explained by the fact that the ductal cells used as a reference are part of the tumor itself, and are exposed to the TME.

Normal lung and skin have squamous components, and consistently we observed no difference in squamous expression in the tumor samples (Fig. 2e). In contrast, the squamous module was induced in the PDAC relative to normal ductal cells, indicating squamous differentiation in the malignant cell population. A similar trend was observed for the basal module (Extended Data Fig. 6f). Several classifications of PDAC have been proposed based on bulk transcriptomics^71–73^, including a distinction between classical (high expression of glandular genes, including *TFF1* and *CEACAM6*) and basal subtypes (high expression of squamous and basal genes, including *LY6D* and *KRT15*). Although squamous cell pancreatic cancer is rare^74,75^, the increase in squamous expression frequency in PDAC suggests that partial metaplasia towards a squamous program is common. Expression of the glandular module was unchanged in LUAD and PDAC relative to their normal counterparts, but increased in SKSC relative to normal skin. This pattern suggests that a malignant population of cells retains expression of modules associated with its cell type of origin (for example, retention of the squamous module in SKSC) and further deploys gene modules from other cell types (increased expression of the glandular module in SKSC).

### Expression of the interferon response is modulated by the tumor microenvironment

Cancer cell states may reflect common physical constraints and interactions with the cellular components of their microenvironment^76–78^. Notably, the success of immunotherapy in a range of cancer types points to conserved interactions between cancer cells and immune cells^19–22^, leading us to ask whether interactions with the immune system shape the set of occurring cancer cell states. The interferon response module in particular may be involved in interactions between cancer cells and the TME. In tumors, type I interferons are secreted by cancer cells and DCs in response to DNA fragments activating the cGAS/STING pathway^79–81^, and result in T cell priming and antitumor activity^44^. IFNγ is mainly produced by adaptive immune cells upon activation^45^, and leads to up-regulation of MHC I genes, initially facilitating tumor rejection but ultimately leading to IFN-unresponsive tumors through immunoediting^82^. Following these observations, we asked whether adaptive immune cells are necessary to elicit the interferon response module in cancer cells *in vivo*. For this, we used an established allograft mouse cancer model in which the TME can be readily perturbed (see Methods). We performed scRNA-Seq on four orthotopic pancreatic tumors to verify that gene modules could be recapitulated in the orthotopic model. Identifying gene modules using NMF as we did previously, we found that five were recapitulated in this system: cycling, stress response, interferon response, hypoxia, and glandular differentiation (Fig. 3a,b, Supplementary Table 4).

**Figure 3:**
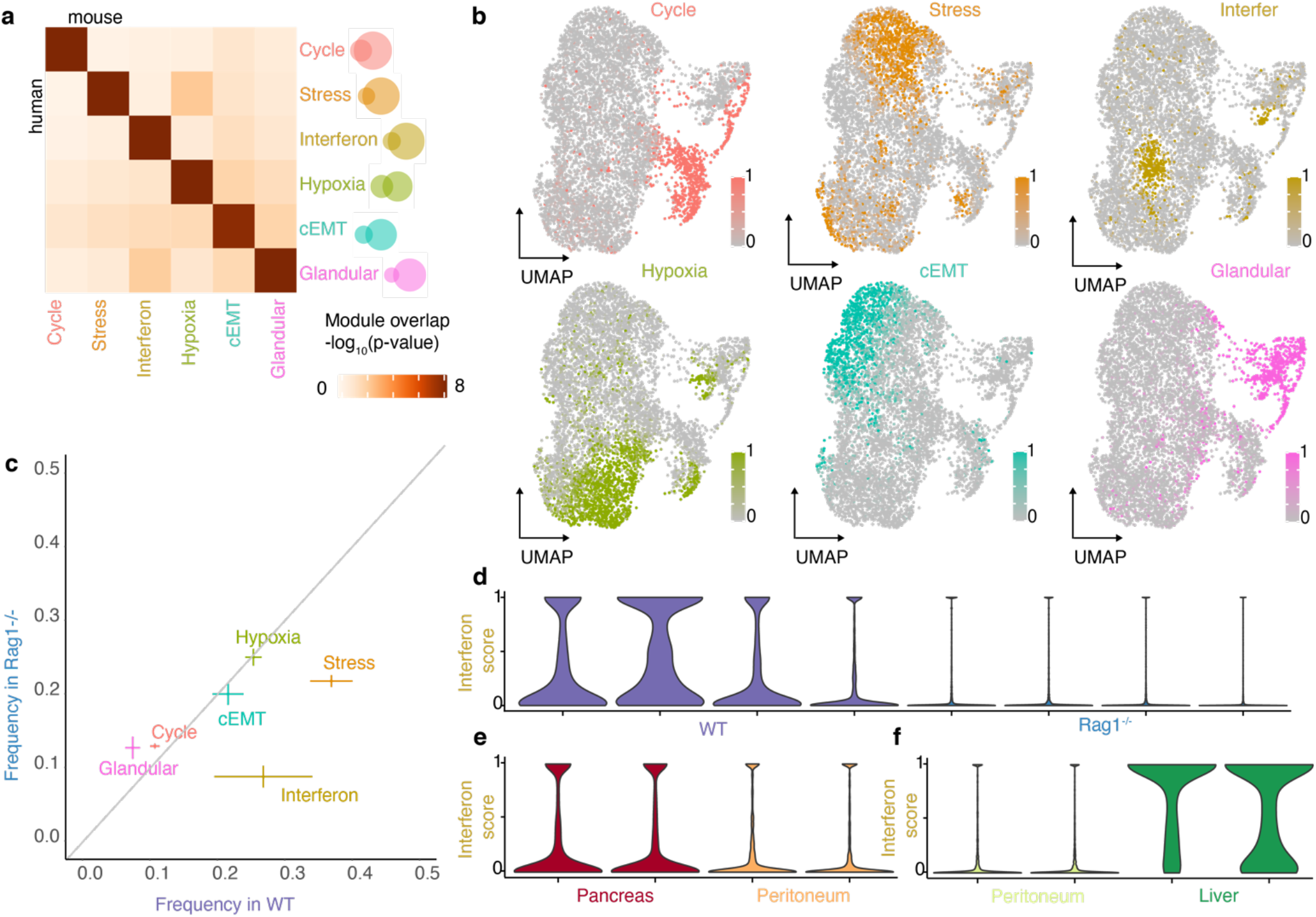
Cancer cell states in perturbed tumor microenvironments. **a**. Heatmap of the significance of the overlap between modules obtained by NMF in an orthotopic model of pancreatic cancer and modules obtained in patient samples in **Figure 1f** (hypergeometric test). **b**. UMAP embedding of malignant cells from orthotopic pancreatic tumors, colored according to the expression score of the six modules shown in **a**. **c**. Module expression frequencies in WT mice vs *Rag1*^-/-^ mice (mean ± standard error). **d**. Violin plots of interferon module expression score in WT mice vs *Rag1*^-/-^ mice. **e**. Same as **d**, for pancreas versus peritoneum. **f**. Same as **d**, for peritoneum versus liver.

In parallel, we collected scRNA-Seq data from four orthotopic tumors formed in *Rag1*^-/-^ mice, which lack T and B cells. Analyzing the gene module expression in the malignant cells from these tumors we found that cycling, stress response, hypoxia and glandular differentiation were expressed at similar frequencies between the *Rag1*^-/-^ and WT mice (Fig. 3c). In contrast, the interferon response module was expressed at lower frequencies in the tumors from the *Rag1*^*-*/-^ mice (*p* < 10^−10^, Kolmogorov-Smirnov test, Fig. 3d). Furthermore, all of the genes of the interferon response module were up-regulated in the interferon response-expressing cells relative to other cancer cells, suggesting that a coordinated response is maintained - albeit in fewer cells (Extended Data Fig. 8d). The MHC I genes of the interferon response module (*B2m, H2-D1, H2-K1*) have a lower overall expression in the *Rag1*^-/-^ mice (although they remain relatively up-regulated in the interferon response-expressing cells), suggesting that lymphocyte depletion has an additional general effect on the expression of MHC I genes which is interferon response-independent.

We next tested whether different tumor microenvironments would also modulate the expression of the interferon response module. In one experiment, we compared the orthotopic tumors in the pancreas, the site of origin of the cancer, to heterotopic tumors in the peritoneum, a common site of metastasis. We found that tumors in the peritoneum have a lower frequency of interferon response-expressing cells (*p* < 10^−6^ in Kolmogorov-Smirnov test, Fig. 3e). In a second experiment, we compared frequencies across two heterotopic sites - peritoneum and liver - in order to model different metastatic sites *in vivo*. Here, we found that the interferon response module is expressed at a higher frequency in the liver relative to the peritoneum (*p* < 10^−10^, Kolmogorov-Smirnov test, Fig. 3f).

Collectively, this set of experiments provides an initial assessment of the occurrence of the interferon response module in cancer. The presence of this module across a variety of cancer types (Fig. 1), organs and immune settings (Fig. 3), suggests that the heterogeneity of interferon response across malignant cells is a common feature of tumors. We found that the adaptive immune system is necessary for most, but not all, of the expression of this module. The remaining expression in the lymphocyte-depleted condition suggests other causes of interferon response in cancer cells, either cancer extrinsic, for example interferon secretion by NK cells, or cancer-intrinsic, consistent with reports of an interferon response module *in vitro*^*83*^. Notably, this finding does not discriminate between signaling mechanisms eliciting an interferon response and long term immunoediting leading to selection of the state within the tumor^40,82^.

### Spatial organization of malignant and non-malignant cell types in the tumor

To further analyze the organization and interactions between cancer cell states and cells of the TME, we turned to sequencing-based spatial transcriptomics (ST)^84^. Unlike scRNA-Seq which is obtained after dissociation – resulting in loss of any spatial information – array-based ST data captures mRNA at each location within the tissue not at single-cell resolution^84^, but rather capturing ∼10 cells per spot. We therefore sought to leverage the properties of both modalities by integrating the paired data for ten tumors (Fig. 4a-c, OVCA NYU1, OVCA NYU3, UCEC NYU3, BRCA NYU0, BRCA NYU1, BRCA NYU2, PDAC NYU1, GIST NYU1, GIST NYU2, LIHC NYU1). Each of our ten tumor ST datasets consisted of ∼2,000 spots (ranging from 1,351 to 2,624) over a 6mm x 6mm area. Spots on the ST array are separated by 100µm allowing us to gain insight into the tumor microenvironment, where, for example, paracrine signaling functions at such distances^85^.

**Figure 4:**
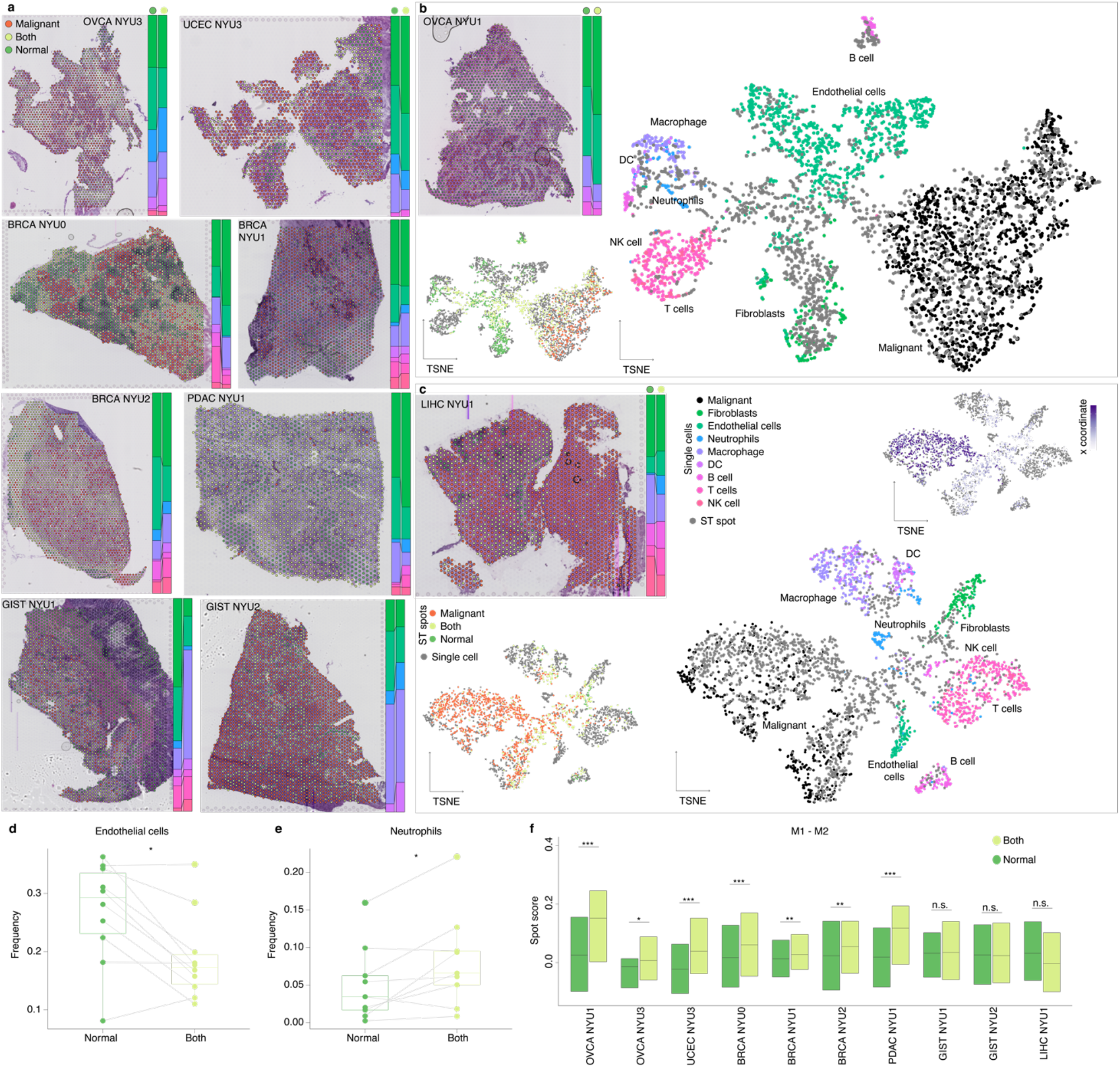
Spatial organization of the tumor microenvironment. **a**. H&E images for the 8 indicated patient tumors overlaid with the locations of the spatial transcriptomic spots colored according to their annotation as ‘Malignant’, ‘Both’, or ‘Normal’. Bar plots indicate the fraction of non-malignant cell types in the ‘Normal’ and ‘Both’ spots for each sample. **b**. Joint dimensionality reduction of single-cell and ST spots for the OVCA NYU1 sample. The top inset indicates the H&E image as in **a**. The bottom inset shows the same joint dimensionality reduction with the ST spots colored according to their annotation. **c**. Joint dimensionality reduction of single-cell and ST spots for the LIHC NYU 1 sample, as in **b**. The top right inset shows the same joint dimensionality reduction with spots colored according to their coordinate along the x axis. **d**. Boxplots of the fractions of endothelial cells and neutrophils in ‘Normal’ and ‘Both’ spots for each sample (*, *p*-value < 0.05; Wilcoxon test). **e**. Boxplots of fractions of neutrophils in ‘Normal’ and ‘Both’ spots for each sample (*, *p-*value < 0.05; Wilcoxon test). **f**. Boxplots of M1-M2 score in ‘Normal’ and ‘Both’ spots for each sample (*, *p*-value < 0.05; ***, *p-*value < 0.001; Wilcoxon test).

Since each spot is a combination of cells of one or more cell types, its transcriptome can be represented as a weighted sum of cell type transcriptomes. To infer the contribution of each cell type at each spot, we performed non-negative linear least squares (NNLS) regression using the average expression profiles of cell types from the paired single cell data (see Methods). We then compared the coefficients obtained for each cell type to those obtained in a random model, and considered a cell to be present in a spot if its coefficient was more than two standard deviations above the mean in the random set. These annotations were further validated by the pathologists on our team (C.H. and D.F.D.). As a framework for further analysis, we divided the spots into three categories according to their cell type annotations: ‘Malignant’, containing only malignant cells, ‘Normal’, containing only immune and stromal cells, and ‘Both’, containing a combination (Fig. 4a, Extended Data Fig. 9, Supplementary File 2).

As an independent method for spot categorization, we also directly compared the spots and single-cell transcriptomes of the 10 paired datasets. Figure 4b shows a dimensional reduction plot of the transcriptomes from the OVCA NYU1 sample, with gray dots indicating the ST spots and the other colors indicating single-cell transcriptomes, colored by the annotated cell types. By invoking mutual nearest neighbor (MNN) integration and joint dimensionality reduction^86^, we found that data from the two modalities are well integrated (see Methods, Fig. 4b,c, Extended Data Fig. 10). The single cells form clusters at the periphery, indicating distinct cell types. The ST spots are either mixed with individual single-cell clusters, indicating a pure population, or bridge multiple clusters, indicating a combination of cell types. Overlaying the spot categories determined by the NNLS method onto this plot, we consistently observed that ‘Malignant’ spots were mixed with the malignant cell cluster, ‘Normal’ spots were in the region of non-malignant cell types, and ‘Both’ spots spanned both malignant and non-malignant single-cell clusters. As a second example, the LIHC ST dataset showed two spatially distinct tumor nodules (Fig. 4c), with the left having substantial mixing between malignant and non-malignant cells and the right consisting of almost only malignant cells. The joint dimensionality reduction analysis reflected the two corresponding malignant clusters, which were not distinct when considering the single-cell dimensionality reduction alone (Extended Data Fig. 12f). This analysis highlights the potential of integrating paired spatial and single-cell datasets to anchor single cells in their spatial context.

To further test the accuracy of the NNLS method to annotate spots, we performed paired scRNA-Seq and ST on two patient-derived melanoma xenografts (PDX) (Extended Data Fig. 11a-b, see Methods). In this setting, only malignant cells are of human origin and therefore express human genes, enabling us to reliably identify malignant cells or spots. Using the NNLS method on the full mouse and human transcriptomes, we first established a ‘ground truth’ for spot identities. We then simulated the patient samples by converting mouse genes to their human orthologs, thereby removing the species information. Annotating the spots using NNLS in this way resulted in 99% and 89% specificity for each sample, supporting its accuracy.

The presence of ‘Normal’ and ‘Both’ spots in each sample enabled us to ask how the cell type composition of the tissue changes in the presence of malignant cells. The fraction of endothelial cells was consistently lower in the spots also containing malignant cells (Fig. 4d), suggesting an incomplete vascularization of the tumor^87,88^. Conversely, neutrophils were found in higher numbers in the ‘Both’ spots (Fig. 4e). Tumor-associated macrophages are broadly defined as M1 - anti-tumor/pro-inflammatory - and M2 - pro-tumor/anti-inflammatory^89–91^. In our single-cell data, we detected two populations of macrophages, one expressing pro-inflammatory genes (e.g., TNF, SPP1, ISG15), and the other characterized by antigen presentation and complement (e.g., HLA-DRA, C1QA, CD163) (Extended Data Fig. 12a,c, Supplementary Table 6). To compare the location of M1 and M2 macrophages relative to that of cancer cells, we scored each spot containing a macrophage for its expression of the signatures of both populations, and calculated the M1-M2 score, which allowed us to compare macrophage polarity across spot categories (Fig. 4f). In the six gynecological samples - ovarian, endometrial and breast cancer - we found that ‘Both’ spots contained a significantly higher M1-M2 score than ‘Normal’ spots, suggesting a robust anti-tumor macrophage activity in proximity to cancer cells. This is in contrast to the findings of a study performed in colorectal carcinoma that detected higher M2 in the tumor relative to adjacent normal tissue^92^. These results suggest the presence of an inflammatory host response surrounding malignant cells within a few 100µms, highlighting the value of studying the architecture of tumors at high resolution. Beyond the M1/M2 dichotomy, macrophage with diverse phenotypes, including pro-angiogenic macrophages^93^ and mesenchymal-like macrophages^94^, are emerging as key players in tumor-immune interactions. A similar analysis for T cell subtypes - cytotoxic, helper and regulatory - did not show consistent results across samples (Extended Data Fig. 12b,d,e, Supplementary Table 7).

### Cancer cell state analysis of tumor cellular neighborhoods

Having identified the malignant and non-malignant cell types within each tumor, we next sought to query the composition of cellular neighborhoods in terms of cancer cell states. For this, we mapped cancer cell states within each ST sample, scoring each ‘Malignant’ spot for its expression of each module (Fig. 5a, see Methods). To establish the validity of this scoring approach, we first turned again to the PDX data and scored the ‘Malignant’ spots for the expression of modules (Extended Data Fig. 11e-f). Since human malignant cells can unambiguously be distinguished from mouse TME cells in this system, we first used the single cell data to confirm that the modules are differentially expressed by malignant cells themselves and rule out the possibility of an artifact stemming from TME contamination. For example, the pEMT module includes genes normally expressed by fibroblasts, but we detected its presence in the malignant cells (Extended Data Fig. 11g-h). The interferon response module was not present, as expected since, like the Rag-/- mice (Fig. 3c), these mice are lymphocyte deficient.

**Figure 5.**
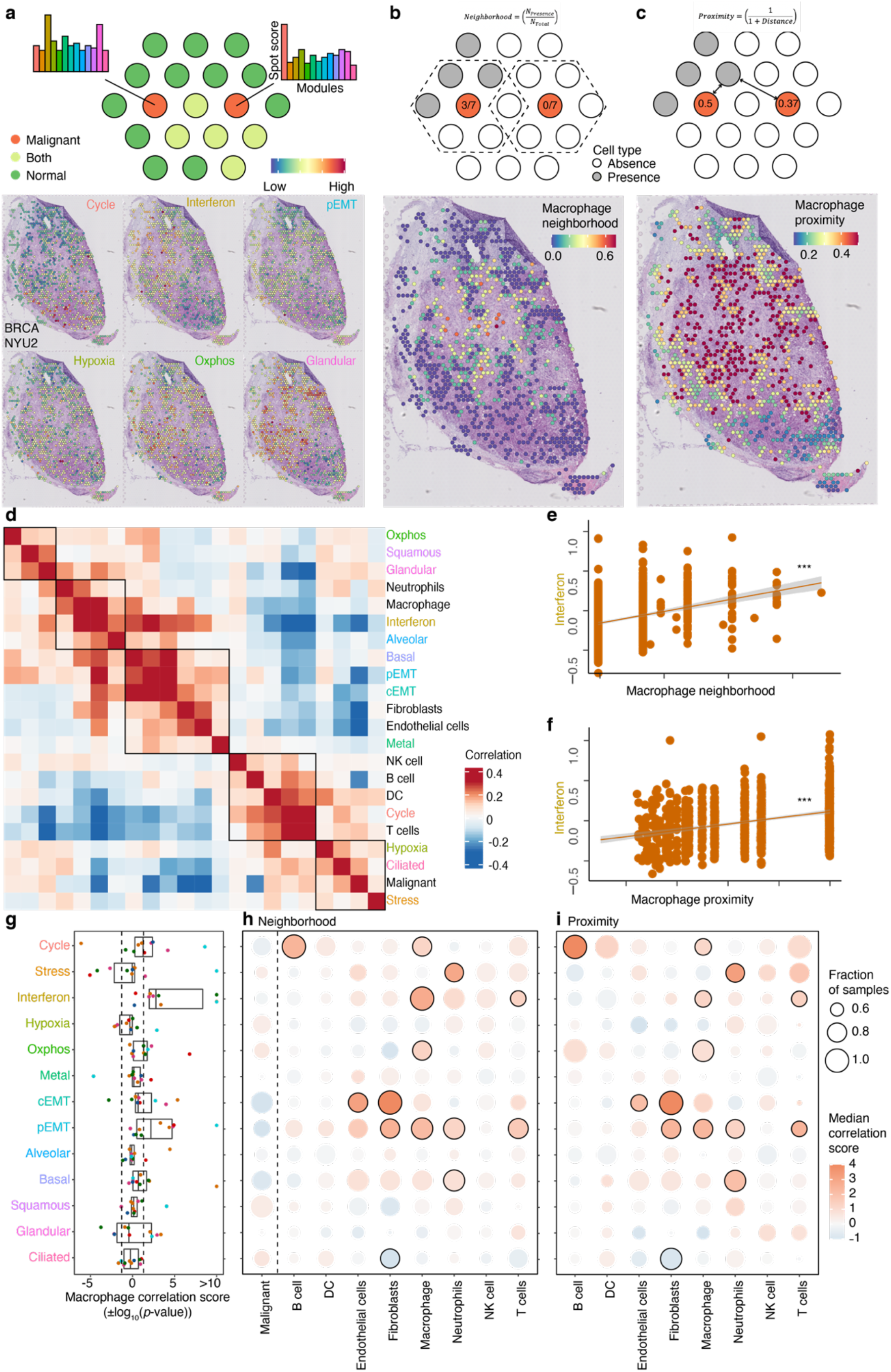
Mapping cancer cell states and their interactions with the TME. **a**. Scoring ST spots for module expression. In the top schematic of spots colored by their annotation as ‘Malignant’, ‘Both’ and ‘Normal’. Only ‘Malignant’ spots are scored for their expression of each module. For the BRCA NYU2 sample, ‘Malignant’ spots are indicated and colored by their expression of the 6 indicated modules. **b**. Characterizing ST spots by cell type neighborhood. In the top schematic grey spots indicate the presence of the cell type of interest, orange indicates ‘Malignant’ spots, and dashed lines their surrounding spots. For the same sample as in **a**, the spots are colored by neighborhood macrophage score. **c**. Characterizing ST spots by cell type proximity. In the top schematic, spots are colored as in b and arrows the distance to the closest spot containing the cell type of interest. For the same sample as in a, the spots are colored by the macrophage proximity score. **d**. Heatmap of correlations between module scores and cell type neighborhood scores for BRCA NYU2 (‘Malignant’ spots only). Boxes indicate clusters of correlated module expression scores and cell type neighborhoods. **e**. Plot of the relationship between the interferon response module score and macrophage neighborhood score in the BRCA NYU2 sample (‘Malignant’ spots only). **f**. Plot of the relationship between the interferon response module score and macrophage proximity in the BRCA NYU2 (‘Malignant’ spots only) **g**. Boxplot of correlation scores (±log_10_(*p*-value)) between module scores and macrophage neighborhood scores across 10 samples, colored as in **Figure 1a**. Positive scores correspond to positive correlations. Dashed lines indicate *p*-value = 0.05. **h**. Correspondence map of significance between module expression scores and cell type neighborhoods. Color represents the median ±log_10_(*p*-value) of the correlation, with red corresponding to positive correlations and blue to negative correlations. Point size represents the fraction of samples in which the correlation is of the same sign as the median correlation. Black outlines indicate relationships where the median ±log_10_(*p*-value) of the correlation was greater than 0.75, and the fraction of samples in which the correlation is of the same sign as the median correlation is greater than 0.5, using both the neighborhood and proximity metrics (see **Fig. 5i**). **i**. Correspondence map of significance between module score and cell type proximity, colored as in **Figure 5h**.

To characterize the cell type composition surrounding each ‘Malignant’ spot, we calculated, for each cell type, two score indices meant to capture their microenvironment. We defined the ‘neighborhood score’ as the fraction of surrounding spots containing that cell type (Fig. 5b, see Methods). This score thus directly measures the cell type composition in the adjacent spots. The proximity score measures how close the spot of interest is to each cell type, and is calculated as the inverse of the shortest distance to a cell of that type (Fig. 5c, see Methods).

Correlating the module scores and cell type neighborhood profiles across the ‘Malignant’ spots of each tumor revealed how cancer cell states and cell types of the TME co-localize to form ‘neighborhoods’ (Fig. 5d). For the BRCA NYU2 tumor shown in Fig. 5a-c, one grouping in the heatmap of relationships contained the interferon response module and macrophages. Studying this correlation more closely confirmed the significance of this positive relationship between the module score and the macrophage neighborhood score (Fig. 5e). A consistent relationship was also observed when computing the presence of macrophages using the proximity score (Fig. 5f). To explore this relationship across all samples, we calculated the correlation score (±log_10_(*p*-value)) of the macrophage neighborhood (Fig. 5g). The correlation with the interferon response was positive in all the samples, and significant for 8 of 10 samples. This was not the case for any other module (Fig. 5g). This suggests that macrophages may elicit the expression of the interferon response module, or that the interferon response-expressing cancer cells may recruit macrophages. Indeed, a recent study showed that stimulation of the interferon response pathway in tumors leads to recruitment and activation of macrophages^95^.

Extending this analysis for all pairs of cell types and modules using both neighborhood and proximity measures (Fig. 5h,i), we identified other consistent co-localizations of cell states with cell types of the TME. In addition to macrophages, the interferon response-expressing cancer cells co-localize with T cells according to both measures (Fig. 5h,i), in line with the finding *in vivo* that lymphocytes lead to increased interferon response (Fig. 3d). Further work is required to establish the mode and directionality of these interactions (Extended Data Fig. 14). Cancer cells undergoing EMT were positively correlated with fibroblasts and endothelial cells, and negatively correlated with other malignant cells, consistent with the finding that they are enriched at the interface of the tumor (Extended Data Fig. 13) and interact with cancer-associated fibroblasts^10^.

To further study the co-localization of macrophages and T cells with interferon response-expressing malignant cells, we turned to CO-Detection by indEXing (CODEX) - a multiplexed protein staining assay - which enables spatial analysis at single-cell resolution^96^. For four of the samples used for spatial transcriptomics (OVCA NYU1, UCEC NYU3, LIHC NYU1, GIST NYU1), we stained for 23 markers (Fig. 6, see Methods), including HLA-DRA as a marker for the interferon response (Supplementary Table 2). Using a supervised gating strategy, we identified malignant cells and cells of the TME, including macrophages, T cells, endothelial cells, and fibroblasts (Extended Data Fig. 15). A subset of PanCytokeratin(PanCK)/EPCAM positive cells was also positive for HLA-DRA, providing evidence at the protein-level that MHCII is differentially expressed in malignant cells of a tumor (Fig. 6a).

**Figure 6:**
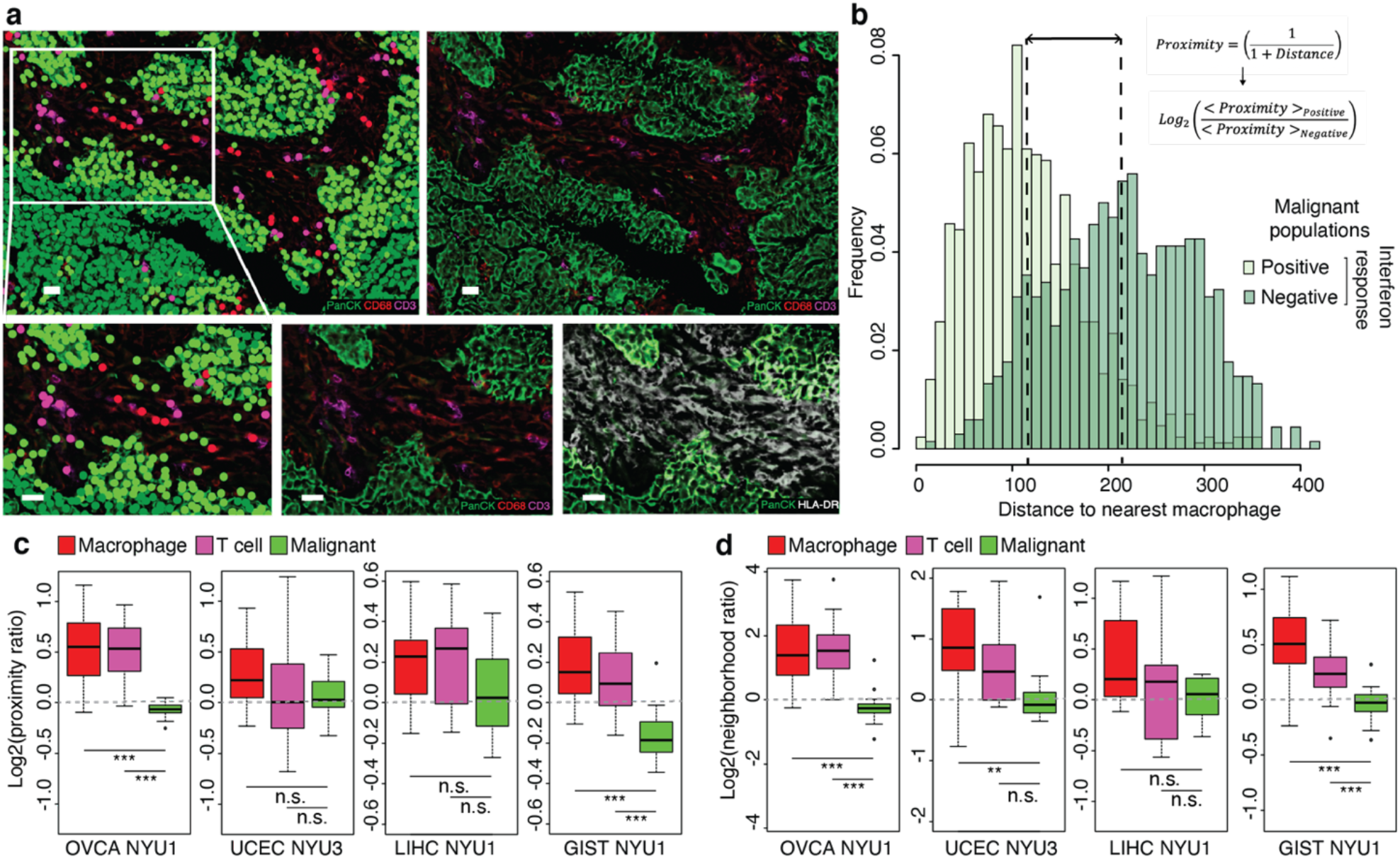
CODEX analysis of samples from four cancer types supporting a proximity of interferon response-expressing malignant cells to macrophages and T cells. **a**. Cell populations and marker expression in a region of OVCA NYU1. Top row displays an entire tile, bottom row displays an enlargement. Top and bottom left: Colored by populations as defined in Extended Data Fig. 15. Top right and bottom center: Colored by expression of markers used to define cell types, as indicated. Bottom right: Colored by expression of PanCK and of HLA-DRA, used to define interferon response positive and negative malignant cells. **b**. For the tile shown in **a**., histogram showing the distance between malignant cells and the nearest macrophage, for interferon-response positive (light green) and negative (dark green) malignant cells. Lines indicate the mean distance for each population, used to calculate the log2(proximity ratio). **c**. Boxplots of the distribution of log2(proximity ratio) of macrophages, T cells and malignant cells across tiles of each sample (*, *p*-value < 0.05; ***, *p-*value < 0.001; two-sided t-test). **d**. Boxplots of the distribution of log2(neighborhood ratio) - see Figure 5b - of macrophages, T cells and malignant cells across tiles of each sample (*, *p*-value < 0.05; ***, *p-*value < 0.001; two-sided t-test).

We therefore defined malignant cells as either interferon response-positive or negative, and compared their proximity to macrophages and T cells. Again, we used the two metrics of proximity and neighborhood (Fig. 5a) to study cell co-localization. For example, for macrophages, the ‘proximity’ of a malignant cell to macrophages is defined as 1/(1+distance), taking the distance to the closest macrophage (Fig. 6b); and the ‘neighborhood’ is defined as the fraction of cells annotated as macrophages within a 100µm radius. For each tile within a given sample, we used the average measure for interferon-positive and for interferon-negative cells to compute a log ratio (Fig. 6b). For both macrophages and T cells, the median log ratio across tiles was positive for both the proximity and neighborhood metrics across the four samples, indicating that interferon response-positive malignant cells are preferentially co-localized with these two cell types relative to interferon response-negative malignant cells (Fig. 6c-d). This suggests that at least a subset of cancer cell states interact with the TME, either being elicited by immune or stromal cells, or altering the cell type composition of their surroundings. Further work will elucidate the causal architecture of these relationships.

## Discussion

Single-cell approaches have greatly advanced our understanding of intra-tumoral heterogeneity, and several studies have led to the demonstration of cancer cell states^6–11,14,36^. While such states were identified in individual cancer types, here we provide the first pan-cancer analysis of transcriptional heterogeneity among malignant cells. Our systematic analysis across cancer types led us to propose a unified catalog of gene modules that underpin recurrent cancer cell states. Building upon findings made in individual cancer types, we identify 16 modules spanning the 15 cancer types studied here - including cycling, stress response, interferon response, hypoxia, and pEMT - as well as modules more specific to cellular identity in specific organ systems - including basal, squamous, glandular, and ciliated differentiation. We expect that future work^97^ will expand the list of cancer cell states and their cancer-type or organ-type specificities as well as pan-cancer features.

Cancer cell states cannot be defined as distinctly as cell types^6,17,36^. Our results indicate that this likely follows from the expression of combinations of modules: since these are not generally mutually exclusive (Fig. 2a, Extended Data Fig. 7), this leads to continuous variation rather than discrete clusters (Fig. 3b). Conversely, distinct states may be observed when the gene modules are mutually exclusive with others, as in the case of the cycling and cilium gene modules (Fig. 2b, Extended Data Fig. 7). Our analysis thus supports the notion that the basic underlying units of tumor transcriptional variability are the gene modules, whose combinatorial products define the cancer cell states. Further work is required to disentangle the relationships that relate the gene modules in terms of their co-expression and mutual exclusivity in determining cell states.

Much of the heterogeneity observed across cancer cells appears to result from redeployment of modules typically expressed in other cellular and developmental contexts^98^. Indeed, our catalog of cancer gene modules includes general features of cell physiology (cycling), specific processes and responses (stress, hypoxia, oxidative phosphorylation, interferon and metal response), and developmental programs (EMT, alveolar, basal, squamous, glandular, ciliated, AC-like, OPC-like, NPC-like). Relative to their normal counterparts, cancer cells exploit the existing gene modules, expressing them at different levels (Fig. 2e, Extended Data Fig. 4d) and more heterogeneously (Extended Data Fig. 4c). The interferon response module, for example, while typically associated with cellular immune response to pathogens, is heterogeneously activated in malignant cells.

It remains unclear to what extent the heterogeneity among cancer cells results from heterogeneity in the signals they receive, or from intrinsic differences between the cells – genetic, epigenetic, or stochastic. Our observations of cancer states across a wide range of cancer types provide evidence that cancer cell states are not genetically defined, but rather represent cellular plasticity. In addition, *in vitro* and *in vivo* studies have shown that cancer cells exhibit a high degree of plasticity and can transition from one state to another^7,83^. In particular, populations seeded by a single state recover the same state proportions as the original tumor^7,83^. Thus, while the individual state identities are highly plastic, their overall distribution may be a stable property. Interestingly, in glioblastoma, tumors harboring different genetic drivers share the same set of states, but differ in the proportions of each state^7^. In this view, an early phase of tumorigenesis would generate the oncogenic background upon which later epigenetic changes would lead to heterogeneity among the malignant population^99^.

Several of the gene modules that we identify here may enable the hallmarks proposed by Hanahan and Weinberg^76,77^, raising the possibility that the cancer hallmarks do not need to be assembled by all individual cells. Rather, cell states may cooperate within the tumor ecosystem leading to higher fitness of the tumor as a whole^78^. For example, induction of angiogenesis or down-regulation of immune surveillance may be mediated by a subset of cancer cells to the benefit of the others. To understand these complex relationships, it is crucial to consider the physical constraints of the tumor, including signaling between neighboring cells, diffusion of oxygen and nutrients, and segregation into niches with distinct composition. The co-localization that we observed of interferon response-expressing cells with T cells and macrophages (Figs. 5,6) highlights that the functional role of cancer cell states may be understood by analyzing the tumor architecture. Recent work has also shown that in glioblastoma macrophages elicit a mesenchymal state among the malignant cells^94^. Furthermore, in head and neck squamous cell carcinoma, pEMT expressing cells were found to be located at the leading edge of the tumor^10^, and this finding appears to be general to several cancer types (Extended Data Fig. 13).

The presence of an interferon response in cancer has been studied extensively, and attempts have been made to harness the response for therapy^100^. Here, we found that genes involved in interferon response are co-regulated and heterogeneously expressed across malignant cells of the tumors in all 15 cancer types studied here (Fig. 1), suggesting that the existence of this state is a necessary feature of tumorigenesis. Indeed, tumors lacking IFNγ receptors fail to develop in mouse models^82^. Paradoxically, however, in tumors containing both interferon responsive and unresponsive cells, the unresponsive cells increase in frequency^82^. Thus, the subset of cells expressing the interferon response module appears to support the growth of other cells within the tumor. This may be explained by the dual function of these genes - with MHCI and MHCII eliciting heightened immune detection of the cells, but PDL1 leading to a generalized increase in immune tolerance^100^.

Understanding cancer cell states has critical implications for therapeutic advances, as intratumoral heterogeneity is a recognized cause of treatment failure and relapse^1–5,101^. In particular, the study of the relationships between cancer cell states and the TME – with an emphasis on immune cell populations – may shed light on the contribution of heterogeneity to tumor fitness, and highlight vulnerabilities that can be exploited for targeted therapy.

## Methods

All of the data from this manuscript has been submitted to GEO with accession number GSE153374.

### Patient tumor scRNA-Seq

#### Data collection and processing

Tumors were collected post-operatively from patients who signed an IRB approved consent to use their biospecimen for research. Each sample was washed in PBS and cut into 4–5-mm^3^ pieces, of which 2-3 were reserved for spatial transcriptomics (see below). The remainder was dissociated for scRNA-Seq using the Miltenyi human tumor dissociation kit according to manufacturer’s instructions. Red blood cell lysis was performed in ACK lysis buffer for 3 minutes. Cells were counted and viability was assessed by trypan blue on a hemocytometer. For samples with low viability (<50%), dead cells were removed using the Miltenyi dead cell separator. Single-cell encapsulation and library preparations were performed using the inDrop platform^102^. Libraries were sequenced on an Illumina NextSeq and reads aligned using a custom inDrop pipeline as previously described^103^. To exclude cells with low quality transcriptomes from analysis, cells with fewer than 500 UMIs or more than 30% mitochondrial or ribosomal reads were filtered out. The Seurat single-cell transformation^86,104^ was used to normalize and center the data, and to identify variable genes.

#### Analysis of previously published data

Published datasets were downloaded from GEO. For PDAC^9^, LIHC^28^, CHCA^29^, LUAD^30^, HNSC^10^, SKSC^31^, THCA^70^, and OGD^14^, raw counts were used and normalized as above. For GBM^7^, normalized data was used and centered, and variable genes were identified using the ‘vst’ method in the Seurat package^86,104^.

#### Cell type annotation and detection of malignant cells

To annotate cell types and identify malignant cells, the following procedure was used.

##### Cell type identification

<u>Part 1:</u> The Seurat package^86^ was used to select variable genes, reduce dimensionality, cluster the cells, and search for differentially expressed genes (using thresholds of p-value < 0.01, percentage of cells expressing > 10%, log fold-change > 0.25, sorted by log fold-change). These genes were cross-referenced with the literature to identify immune (expressing *PTPRC, CD19, CD4, CD8A, FOXP3, CD68, S100A8, MS4A2*), stromal (expressing *HBA1, PECAM1, COL4A1* or *COL1A1*), and epithelial (*EPCAM, KRT*) cell types. <u>Part 2:</u> The SingleR package^25^ was used with the Human Primary Cell Atlas database^105^ to annotate each cell as a cell type. Cells that received the following SingleR annotations were classified as non-malignant: B_cell, BM, BM & Prog., CMP, DC, Endothelial_cells Erythroblast, Gametocytes, GMP, HSC_-G-CSF, HSC_CD34+, Macrophage, MEP, Monocyte, MSC, Myelocyte, Neutrophils, NK_cell, Osteoblasts, Platelets, Pre-B_cell_CD34-, Pro-B_cell_CD34+, Pro-Myelocyte, and T_cells. Differential gene expression was performed on each of the identified cell types (Extended Data Fig.1c, Supplementary File 1). These annotations served as the basis for annotation of the spatial transcriptomic spots, and were used to validate the annotation of clusters identified by gene expression in Part 1.

##### Malignant cell identification

<u>Part 1:</u> For epithelial and stromal tumors, the expression pattern of the epithelial and stromal cluster respectively was examined to distinguish malignant from non-malignant cells (Supplementary File 1). Specifically, genes were identified which are overexpressed in malignant relative to normal tissue for each cancer type: *WFDC2*^*106*^ for OVCA and UCEC; *CLU*^*107*^ and *MGP*^*107*^ for BRCA; *LAMC2*^*108*^and *TM4SF1*^109^ for PDAC; *CEACAM5*^*110*^ for COAD; *APOH*^*111*^ for LIHC; *TMPRSS2*^*112*^ and *CLDN4*^*112*^ for PRAD; *CA9*^*113*^ for KIRC; *PDGFRA, KCNK3*^*114*^ and *ANO1*^*115*^ for GIST. <u>Part 2:</u> RNA-based copy-number variation inference was performed on the putative set of malignant cells, as implemented in the inferCNV package^26^, using all other cells from the sample as a reference and searching for consistent patterns of copy-number variation (Extended Data Fig. 2b). <u>Part 3:</u> Dimensionality reduction was performed on the putative sets of malignant cells from different tumors to validate that they form separate clusters, a known property of malignant cells^36^ (Extended Data Fig. 2a).

#### Non-negative matrix factorization (NMF) and module detection

NMF was performed separately on the identified malignant cells of each sample (Fig. 1c). Starting from the normalized centered expression of variable genes, all negative values were set to 0, as previously described^10^. The “nsNMF” method was applied for ranks between 5 and 25 – as implemented in the NMF R package^10,116^. To define non-overlapping gene modules, a previously described gene ranking algorithm was implemented^117^. Beginning with the matrix of the contribution of genes (rows) to the factors (columns), two ranking matrices were constructed, (list 1) ranking the gene contributions to each factor and (list 2) ranking for each gene the factors to which it contributes. For each factor, genes were added in the order of their contribution (list 1), until a gene was reached which contributed more to another factor, i.e. its rank across factors (list 2) was not 1. Factors which yielded fewer than 5 genes were removed, and the procedure repeated. With this method, the number of modules was at most the rank of the NMF, and the modules were robust to the rank chosen. The highest rank for which the number of modules was equal to the rank was selected for downstream analysis.

#### Graph-based clustering and identification of consensus gene modules

The full list of modules obtained for individual tumors was filtered to retain only those with at least 5% overlap (by Jaccard index) with at least 2 other modules. An adjacency matrix was then constructed connecting genes according to the number of individual tumor modules in which they co-occur. Gene-gene connections were filtered out if they occurred in fewer than 2 individual tumor modules, and genes with fewer than 3 connections were removed. The graph was clustered using infomap clustering implemented in the igraph package^118^. Finally, modules with potential biological relevance were retained by filtering out those with fewer than 5 genes or without significant overlap with gene ontology terms. The final graph (Extended Data Fig. 3f) was visualized with the fruchterman-reingold layout.

#### SCENIC module identification and module comparison

SCENIC regulon identification was performed using the SCENIC package^69^ implemented in R and Python. Genes were filtered to have at least 0.05 counts per cell on average and to be detected in at least 5% of the cells, and the transcription factor-binding databases used were 500bp-upstream and tss-centered-10kb. To compare modules obtained by NMF in individual tumors to each other (Fig. 1e, Extended Data Fig. 3g), the significance of the pairwise overlap was calculated using the hypergeometric distribution. SCENIC-derived regulons were similarly compared to the consensus modules (Extended Data Fig. 3h), and were considered to match a consensus module if the *p*-value of the overlap was <10^−3^. Transcription factors annotated to regulons matching each consensus module were then tabulated (Supplementary Table 4) and the top 1-2 factors found in multiple samples are shown in Figure 1f ‘Regulators’.

#### Module annotation and receptor-ligand analysis

Gene Ontology terms were accessed using the MSigDB package in R^119^ (Extended Data Fig. 2a). Cell type markers were downloaded from PanglaoDB^120^ (Extended Data Fig. 2b). Tumor-derived signatures were accessed from Neftel et al.^7^ (Extended Data Fig. 2c)^,,^ Puram et al.^10^ (Extended Data Fig. 2d) and Ji et al.^31^ (Extended Data Fig. 2e) The significance of the overlap between each consensus module and each downloaded gene set was calculated using the hypergeometric distribution. Receptor-ligand analysis (Extended Data Fig. 14) was performed using Nichenet.

#### Significance of module presence

The previously described ‘overdispersion’ approach was used to quantify the differential expression of a particular module in a set of malignant cells (Extended Data Fig. 4a-c)^32^. For each module, PCA was performed on the expression of the module genes, and the variance explained by PC1 was calculated. For each module, 10^3^ random lists of genes with similar expression levels were generated as has been done previously^6^, and the variance explained by PC1 in those genesets was calculated. The significance of the presence was calculated as -log_10_(*p*), where *p* is the fraction of random genesets that resulted in a higher PC1 variance than the module itself. This enabled the identification of tumors in which specific modules are differentially expressed in a statistically significant manner.

#### Module expression scoring

The expression level of each module in individual cells (Fig. 2a,d, Extended Data Fig. 6b,d) was scored as follows. For each module, 10^3^ random lists of genes with similar expression levels were generated as has been done previously^6^. For each cell, the average centered expression of these genesets was calculated, along with that of the module genes. *p* was defined as the fraction of random genesets with a higher average expression than the module itself. The score was defined as -log_10_(*p*) and rescaled linearly to [0,1]. A module was considered expressed in a given cell if its score was higher than 0.5, and these binary values were used to calculate the frequency of expression of each module in each sample (Fig. 2e, Extended Data Fig. 6e-g). The matrix of module expression scores was used to perform TSNE with 500 iterations and a perplexity of 100 (Fig. 2b-d, Extended Data Fig. 6a-c). The mixing of tumors in the TSNE was assessed by calculating the entropy at each point using its 20 nearest neighbors (Extended Data Fig. 6c).

#### Analysis of normal epithelial cells

For normal fallopian tube^33^, breast^34^ epithelium and normal liver^35^, single-cell RNA-Seq data was downloaded from GEO. For LUAD^30^ and SKSC^31^, single-cell RNA-Seq of matched samples representing normal lung and skin were downloaded from GEO as for the tumor samples. Cells were then annotated according to the lines of evidence 1. and 2. (see ‘Cell type annotation and detection of malignant cells’) to identify epithelial cells. For pancreas, the single-cell RNA-Seq data collected from PDAC^9^ contained malignant as well as non-malignant epithelial cells, with non-malignant cells expressing high levels of epithelial cell markers (e.g., *EPCAM*) but low levels of cancer-specific genes (e.g., *LAMC2, CDKN2A* and *TM4SF1*) and displaying low CNV^9^. Further analysis of each normal cell dataset, including significance of module presence and scoring module expression, was performed as for the malignant cell datasets.

#### TCGA survival analysis

Scoring and survival analysis were performed as in Cook et al.^68^. Expression profiles were obtained from https://gdc.cancer.gov/node/905, normalized and z-scored. To infer the expression of modules in these bulk RNA-Seq samples, modules were first filtered by calculating the specificity of each gene for each cell type using the genesorteR package^121^ (Extended Data Fig. 16a), and retaining genes whose median specificity across tumor samples was highest in malignant cells (Extended Data Fig. 16b). Samples were then scored by calculating the average z-score expression of these filtered modules. The proportions of leukocytes and stromal cells were accessed from Thorsson et al.^122^. Association with progression free survival was calculating using a Cox proportional hazards model with the following independent covariates: age, gender, stage, leukocyte fraction, stromal fraction, and expression of cycle, stress, interferon, pEMT, basal, squamous and glandular modules (Extended Data Fig. 16c).

### Patient tumor spatial transcriptomics

#### Data collection and processing

From the 10 tumors (OVCA NYU1, OVCA NYU3, UCEC NYU3, BRCA NYU0, BRCA NYU1, BRCA NYU2, PDAC NYU1, GIST NYU1, GIST NYU2, LIHC NYU1), 2-3 pieces were embedded in OCT by placing them cut side down into a plastic mold. The OCT-filled mold was then snap frozen in chilled isopentane and stored at -80°C until use. Cryosections were then cut at 10µm thickness and mounted onto Visium arrays. Tissue optimization and library preparation were performed according to manufacturer’s instructions, with 12 minutes of permeabilization. Libraries were sequenced on an Illumina NextSeq and aligned using the Visium SpaceRanger pipeline. As a quality control step, spots with fewer than 500 UMIs or more than 30% mitochondrial or ribosomal reads were filtered out. The Seurat single-cell transformation ^86,104^ was used to normalize and center the data, and to identify variable genes.

#### Deconvolution of spatial transcriptomic spots

Spots were annotated using three parallel methods. First, non-negative least squares (NNLS) regression was performed using the single-cell RNA-Seq expression profiles. Specifically, average profiles were calculated for each cell type (annotated using the SingleR package, see ‘Cell type annotation and detection of malignant cells’), using only the paired sample when possible (i.e. when at least 20 cells of that type were present) or the pooled expression profiles from all samples. These profiles were then used to perform linear regression on each spot using the NNLS package in R^123^ and obtain estimates for the coefficient of each cell type at each spot (Fig. 4a-c, Extended Data Fig. 4). The genes used were the intersection of variable genes in the single cell data and spatially variable genes in the spatial transcriptomic data, obtained with ‘FindVariableFeatures’ and ‘FindSpatiallyVariableFeatures’ respectively^86,123^. Because the distributions of regression coefficients varied across cell types, and were not usually bimodal, thresholds for cell type presence/absence were set for each cell type individually using a null distribution of coefficients in the sample, as follows. First, spots were selected which had a predicted score of 0 for the cell type in question (see below for mutual nearest neighbor annotation prediction). The resulting gene expression matrix was shuffled and used for NNLS, in 100 independent iterations. The distribution of coefficients for the cell type in question was then used to set the threshold at the mean + 2 x standard deviations. Second, mutual nearest neighbor (MNN) integration was performed using the Seurat package^86,123^, using the same set of genes as for NNLS. Prediction coefficients were obtained using the ‘TransferData’ function, and were binarized using a threshold of 0.9. Finally, NMF was performed on each dataset as described for the scRNA-Seq data (see ‘NMF’). The output was processed as above (see ‘NMF’), and factors were named according to the gene with the highest coefficient (Extended Data Fig. 4).

#### Annotation of spatial transcriptomic spots

Signatures for M1 and M2 macrophages and for cytotoxic, helper and regulatory T cells were obtained by performing differential gene expression on the macrophage population of OVCA NYU1, keeping the top 100 genes by *p*-value for each cluster (Extended Data Fig. 12a-b). The inflammatory and myofibroblast signatures were downloaded from Elyada et al.^124^. To ensure that the increase in M1-M2 score in the ‘Both’ spots relative to ‘Normal’ was not due to the presence of malignant cells themselves, we also scores the single-cell RNA-Seq data for the signatures, and confirmed that macrophages were the only cell type with wide bimodal distribution of M1-M2 scores (Extended Data Fig. 12c). Spatial transcriptomic spots containing macrophages, T cells or fibroblasts were scored for the expression of the respective signatures using the ‘AddModuleScore’ function in Seurat^86,123^ (Fig. 4f). Distances were calculated using euclidean distance on the pixel coordinates, and scaled such that the unit is inter-spot distance (100µm). The depth of malignant spots was calculated as the shortest distance to a spot containing a non-malignant cell type. Proximity was defined as 1/(1+distance) (Fig. 5c). The neighborhood of a spot was defined as spots of distance ≤ 1 (including the spot itself), resulting in sets of ≤ 7 neighbors per spot. The neighborhood cell type fraction was then calculated from the binarized cell type annotations of this set (Fig. 5b). ‘Malignant’ spots were scored for the expression of each module using the Seurat function “AddModuleScore” (Fig. 5a)^86,123^.

### Patient tumor CODEX

#### Staining and image acquisition

Four fresh frozen samples (OVCA NYU1, UCEC NYU3, LIHC NYU1, GIST NYU1) were cryosectioned at 10µm thickness and mounted onto a glass coverslip coated with poly-lysine. The tissue was stained and the CODEX multicycle reaction performed as described in the CODEX user manual Revision C. Briefly, the sample coverslip was placed on Drierite beads for 5 minutes, then incubated in Acetone for 10 minutes and set in a humidity chamber for 2 minutes. The sample was hydrated in CODEX Hydration Buffer for twice for 2 minutes, fixed in 1.6% Paraformaldehyde in CODEX Hydration Buffer for 10 minutes, then washed in CODEX Hydration Buffer twice for 2 minutes. The sample was then equilibrated in CODEX Staining Buffer for 30 minutes, then stained with a barcoded antibody cocktail in CODEX Blocking Buffer for 3 hours in a humidity chamber. Antibodies comprising the antibody cocktail are listed in Table 7. The sample was washed three times for 2 minutes in CODEX Staining Buffer, fixed in 1.6% Paraformaldehyde in CODEX Storage Buffer for 10 minutes, and then washed 3 times in 1X PBS. The sample was incubated in 4°C methanol for 5 minutes, washed 3 times in 1X PBS, and then fixed using the CODEX Final Fixative Reagent Solution for 20 minutes in a humidity chamber. The sample was washed three times in PBS and then stored in CODEX Storage Buffer for 3 days until imaging. A commercial Akoya CODEX instrument and a Keyence BZ-X800 microscope with a 20x Nikon PlanApo NA 0.75 objective were used to treat and image tissue using complementary fluorescent reporters. The protocols from Akoya CODEX user manual revision C were followed. The four samples were imaged and processed in one CODEX run. Every sample was imaged in 64 (8 × 8) tiles.

#### Image processing

Raw TIFF image files were processed using the CODEX Processor. ImageJ and its CODEX Multiplex Analysis Viewer (MAV) plugin were used to visualize, annotate and define cell populations. Supervised clustering was used to define populations for each tissue. Briefly, the gating function in CodexMAV was used to define cells as positive by setting a gate on log10 intensity vs frequency of the marker of interest, for the non-malignant cell populations. To define malignant cell populations in epithelial cancers log10 EpCAM versus log10 PanCK intensities were used to define a double positive population. For the GI stromal tumor, log10 Podoplanin vs. αSMA intensities were used, where double positive cells were annotated as muscle and Podoplanin+ αSMA-were annotated as malignant. Further characterization of the malignant interferon response cell state of the malignant cells, in both tumor types, was conducted by gating on the log10 HLA-DR intensity vs. frequency. All gating was performed on the segmented image of each sample separately, and was based on marker intensity value independent of the visualized image. The threshold dictating the gate was verified visually by validating overlap between the defined population and marker expression, and was set for each marker and for each sample independently. Gating for each population was done sequentially to avoid overlap between populations. The x and y coordinates and the annotation of each cell were exported for further analysis. Spatial analysis was done in R for each sample independently. For each tile of the image, a distance matrix was calculated and used to quantify the distribution of distances closest to populations of interest (e.g Macrophage) in interferon and non-interferon malignant cells, or the distribution of number of cells of interest in a distance of choice. Tiles with obvious bubbles, tissue folding issues, non-malignant cell dominance (more than 70%, mainly the blood vessel areas), or not containing any tissue, were excluded from the analysis.

### Orthotopic pancreatic tumor mouse models and scRNA-Seq

#### Tumor collection

All experiments were approved by the New York University School of Medicine Institutional Animal Care and Use Committee. *Rag1*^*-/-*^ and WT C57BL/6 mice were obtained from Jackson Laboratories (Bar Harbor, ME). The *Kras*^*G12D*^*;Tp53*^*R172H*^*;Pdx1*^*Cre*^ (KPC) derived cell line FC1242 was utilized for orthotopic injection of 100,000 cells into the tail of pancreata of 8-12 week old C57BL/6 or *Rag*^*-/-*^ mice. To model liver and peritoneal metastases, mice received FC1242 via splenic (10^6^ cells) and intraperitoneal (10^5^ cells) injection, respectively. Tumors were harvested 2-3 weeks after injection and dissociated using Miltenyi mouse tumor dissociation kit enzymes D and R according to manufacturer’s instructions. Red blood cell lysis was performed for 3 minutes in ACK lysis buffer. Dead cells were removed using the Miltenyi dead cell separator. In order to hash and pool replicates, cells were then labeled with Biolegend oligonucleotide-conjugated antibodies according to manufacturer’s instructions. Single-cell encapsulation and library preparation were performed using the 10x Genomics Chromium. Libraries were sequenced on an Illumina NextSeq and reads aligned using the 10x Genomics CellRanger pipeline.

#### Transcriptomic analysis of mouse scRNA-Seq data

Quality control and processing were performed separately for each mouse scRNA-Seq sample separately as for the human data (‘Patient tumor scRNA-Seq’). Samples from all 3 experiments were then combined for cell type annotation. NMF and module identification was performed for the 4 pancreatic WT tumors together. These modules were compared to the consensus modules obtained from patient tumors by orthology mapping using the biomart database^125^. Overlap between modules across species was tested using the hypergeometric distribution. Module expression was scored for each experiment separately as for the human data, and frequencies were compared across conditions using Kolmogorov-Smirnov tests on the distributions (maximum *p*-value of pairwise comparisons across conditions is reported).

### Patient derived melanoma xenograft (PDX) models and scRNA-Seq

#### Tumors collection

Samples were obtained from the Hernando lab. Briefly, NSG (Jax 005557) mice were obtained from Jackson Laboratories (Bar Harbor, ME). Cells obtained from patient melanoma brain metastases were injected intradermally and collected after 82 days. For single-cell RNA-Seq, the sample was minced and incubated in 4mL HBSS buffer with 1mg Collagenase IV and 12.5uL DNAse I for 10min at 37C. Single-cell encapsulation and library preparation were performed using the 10x Genomics Chromium as above (‘Orthotopic pancreatic tumor mouse models and scRNA-Seq’). For spatial transcriptomics, samples were embedded in OCT and processed by Visium according to manufacturer’s instructions as above (‘Patient tumor spatial transcriptomics’). Libraries were sequenced on an Illumina NextSeq and reads aligned using the 10x Genomics pipelines.

#### Transcriptomic analysis of scRNA-Seq PDX data

Quality control and processing were performed as for the human data (‘Patient tumor scRNA-Seq’). Cells with nCount_human > 100 x nCount_mouse were annotated as human, cells with nCount_mouse > nCount_human were annotated as mouse, and other cells were discarded. The presence of modules in human cells was calculated as above (‘Significance of module presence’). Mouse cells were further clustered using Seurat and annotated using marker genes to obtain cell types for spatial transcriptomic deconvolution.

#### Transcriptomic analysis of ST PDX data

Spatial transcriptomics analysis was performed as for the human data (‘Patient tumor spatial transcriptomics’). For NNLS deconvolution, the expression of human and mouse genes was used to generate a ‘ground truth’, and human orthologs of mouse genes were used to simulate patient tumors (Extended Data Fig. 11e-f).

## Acknowledgements

We thank Rich White, Felicia Kuperwaser, Rahul Satija, and Benjamin Neel for critical readings and helpful suggestions. We thank Eva Hernando for helpful discussions and for providing the PDX samples. We thank the Center for Biospecimen Research and Development at NYU Langone Health. The authors declare that they have no conflict of interest relating to this work.

## Notes

### Competing Interest Statement

The authors have declared no competing interest.

## REFERENCES

1. Easwaran, H., Tsai, H.-C. & Baylin, S. B. Cancer epigenetics: tumor heterogeneity, plasticity of stem-like states, and drug resistance. Mol. Cell 54, 716–727 (2014).

2. McGranahan, N. & Swanton, C. Clonal Heterogeneity and Tumor Evolution: Past, Present, and the Future. Cell 168, 613–628 (2017).

3. Marusyk, A. & Polyak, K. Tumor heterogeneity: Causes and consequences. Biochimica et Biophysica Acta (BBA) - Reviews on Cancer vol. 1805 105–117 (2010).

4. Heppner, G. H. & Miller, B. E. Tumor heterogeneity: biological implications and therapeutic consequences. Cancer Metastasis Rev. 2, 5–23 (1983).

5. Alizadeh, A. A. et al. Toward understanding and exploiting tumor heterogeneity. Nat. Med. 21, 846–853 (2015).

6. Patel, A. P. et al. Single-cell RNA-seq highlights intratumoral heterogeneity in primary glioblastoma. Science 344, 1396–1401 (2014).

7. Neftel, C. et al. An Integrative Model of Cellular States, Plasticity, and Genetics for Glioblastoma. Cell 178, 835–849.e21 (2019).

8. Baron, M. et al. A population of stress-like cancer cells in melanoma promotes tumorigenesis and confers drug resistance. doi:10.1101/396622.

9. Moncada, R. et al. Integrating microarray-based spatial transcriptomics and single-cell RNA-seq reveals tissue architecture in pancreatic ductal adenocarcinomas. Nat. Biotechnol. 38, 333–342 (2020).

10. Puram, S. V. et al. Single-Cell Transcriptomic Analysis of Primary and Metastatic Tumor Ecosystems in Head and Neck Cancer. Cell 171, 1611–1624.e24 (2017).

11. Kim, C. et al. Chemoresistance Evolution in Triple-Negative Breast Cancer Delineated by Single-Cell Sequencing. Cell 173, 879–893.e13 (2018).

12. Izar, B. et al. A single-cell landscape of high-grade serous ovarian cancer. Nat. Med. (2020) doi:10.1038/s41591-020-0926-0.

13. Barkley, D., Rao, A., Pour, M., França, G. S. & Yanai, I. Cancer cell states and emergent properties of the dynamic tumor system. Genome Res. 31, 1719–1727 (2021).

14. Tirosh, I. et al. Single-cell RNA-seq supports a developmental hierarchy in human oligodendroglioma. Nature 539, 309–313 (2016).

15. Reitman, Z. J. et al. Mitogenic and progenitor gene programmes in single pilocytic astrocytoma cells. Nat. Commun. 10, 3731 (2019).

16. Rambow, F. et al. Toward Minimal Residual Disease-Directed Therapy in Melanoma. Cell 174, 843–855.e19 (2018).

17. Baron, M. et al. The Stress-Like Cancer Cell State Is a Consistent Component of Tumorigenesis. Cell Syst 11, 536–546.e7 (2020).

18. Siemann, D. W. Tumor Microenvironment. (John Wiley & Sons, 2011).

19. Mellman, I., Coukos, G. & Dranoff, G. Cancer immunotherapy comes of age. Nature vol. 480 480–489 (2011).

20. Ribas, A. & Wolchok, J. D. Cancer immunotherapy using checkpoint blockade. Science vol. 359 1350–1355 (2018).

21. Chuang, Y.-C. et al. Adjuvant Effect of Toll-Like Receptor 9 Activation on Cancer Immunotherapy Using Checkpoint Blockade. Front. Immunol. 11, 1075 (2020).

22. Waldmann, T. A. Immunotherapy: past, present and future. Nature Medicine vol. 9 269–277 (2003).

23. Dirkse, A. et al. Stem cell-associated heterogeneity in Glioblastoma results from intrinsic tumor plasticity shaped by the microenvironment. Nat. Commun. 10, 1787 (2019).

24. Cazet, A. S. et al. Targeting stromal remodeling and cancer stem cell plasticity overcomes chemoresistance in triple negative breast cancer. Nat. Commun. 9, 2897 (2018).

25. Aran, D. et al. Reference-based analysis of lung single-cell sequencing reveals a transitional profibrotic macrophage. Nat. Immunol. 20, 163–172 (2019).

26. Website. inferCNV of the Trinity CTAT Project. https://github.com/broadinstitute/inferCNV.

27. Lin, W. et al. Single-cell transcriptome analysis of tumor and stromal compartments of pancreatic ductal adenocarcinoma primary tumors and metastatic lesions. Genome Med. 12, 80 (2020).

28. Sharma, A. et al. Onco-fetal Reprogramming of Endothelial Cells Drives Immunosuppressive Macrophages in Hepatocellular Carcinoma. Cell 183, 377–394.e21 (2020).

29. Zhang, M. et al. Single-cell transcriptomic architecture and intercellular crosstalk of human intrahepatic cholangiocarcinoma. J. Hepatol. 73, 1118–1130 (2020).

30. Kim, N. et al. Single-cell RNA sequencing demonstrates the molecular and cellular reprogramming of metastatic lung adenocarcinoma. Nat. Commun. 11, 2285 (2020).

31. Ji, A. L. et al. Multimodal Analysis of Composition and Spatial Architecture in Human Squamous Cell Carcinoma. Cell 182, 1661–1662 (2020).

32. Cantini, L. et al. Classification of gene signatures for their information value and functional redundancy. NPJ Syst Biol Appl 4, 2 (2018).

33. Hu, Z. et al. The Repertoire of Serous Ovarian Cancer Non-genetic Heterogeneity Revealed by Single-Cell Sequencing of Normal Fallopian Tube Epithelial Cells. Cancer Cell 37, 226–242.e7 (2020).

34. Nguyen, Q. H. et al. Profiling human breast epithelial cells using single cell RNA sequencing identifies cell diversity. Nat. Commun. 9, 2028 (2018).

35. MacParland, S. A. et al. Single cell RNA sequencing of human liver reveals distinct intrahepatic macrophage populations. Nat. Commun. 9, 4383 (2018).

36. Tirosh, I. et al. Dissecting the multicellular ecosystem of metastatic melanoma by single-cell RNA-seq. Science 352, 189–196 (2016).

37. Brady, S. W. et al. Combating subclonal evolution of resistant cancer phenotypes. Nat. Commun. 8, 1231 (2017).

38. Vilgelm, A. E. & Richmond, A. Chemokines Modulate Immune Surveillance in Tumorigenesis, Metastasis, and Response to Immunotherapy. Frontiers in Immunology vol. 10 (2019).

39. Wan, S. et al. Chemotherapeutics and radiation stimulate MHC class I expression through elevated interferon-beta signaling in breast cancer cells. PLoS One 7, e32542 (2012).

40. Dunn, G. P., Koebel, C. M. & Schreiber, R. D. Interferons, immunity and cancer immunoediting. Nat. Rev. Immunol. 6, 836–848 (2006).

41. Park, I. A. et al. Expression of the MHC class II in triple-negative breast cancer is associated with tumor-infiltrating lymphocytes and interferon signaling. PLOS ONE vol. 12 e0182786 (2017).

42. Axelrod, M. L., Cook, R. S., Johnson, D. B. & Balko, J. M. Biological Consequences of MHC-II Expression by Tumor Cells in Cancer. Clin. Cancer Res. 25, 2392–2402 (2019).

43. Borden, E. C. Interferons α and β in cancer: therapeutic opportunities from new insights. Nat. Rev. Drug Discov. 18, 219–234 (2019).

44. Dunn, G. P. et al. A critical function for type I interferons in cancer immunoediting. Nat. Immunol. 6, 722–729 (2005).

45. Parker, B. S., Rautela, J. & Hertzog, P. J. Antitumour actions of interferons: implications for cancer therapy. Nat. Rev. Cancer 16, 131–144 (2016).

46. Kinker, G. S. et al. Pan-cancer single-cell RNA-seq identifies recurring programs of cellular heterogeneity. Nat. Genet. 52, 1208–1218 (2020).

47. Jögi, A. Tumour Hypoxia and the Hypoxia-Inducible Transcription Factors: Key Players in Cancer Progression and Metastasis. Tumor Cell Metabolism 65–98 (2015) doi:10.1007/978-3-7091-1824-5_4.

48. Eales, K. L., Hollinshead, K. E. R. & Tennant, D. A. Hypoxia and metabolic adaptation of cancer cells. Oncogenesis vol. 5 e190–e190 (2016).

49. Al Tameemi, W., Dale, T. P., Al-Jumaily, R. M. K. & Forsyth, N. R. Hypoxia-Modified Cancer Cell Metabolism. Front Cell Dev Biol 7, 4 (2019).

50. Sethi, N. et al. Mutant p53 induces a hypoxia transcriptional program in gastric and esophageal adenocarcinoma. JCI Insight 4, (2019).

51. Rankin, E. B. & Giaccia, A. J. Hypoxic control of metastasis. Science vol. 352 175–180 (2016).

52. Semenza, G. L. Hypoxia-inducible factors: mediators of cancer progression and targets for cancer therapy. Trends Pharmacol. Sci. 33, 207–214 (2012).

53. Ashton, T. M., McKenna, W. G., Kunz-Schughart, L. A. & Higgins, G. S. Oxidative Phosphorylation as an Emerging Target in Cancer Therapy. Clin. Cancer Res. 24, 2482–2490 (2018).

54. Zheng, J. Energy metabolism of cancer: Glycolysis versus oxidative phosphorylation (Review). Oncol. Lett. 4, 1151–1157 (2012).

55. Cherian, M. G., Jayasurya, A. & Bay, B.-H. Metallothioneins in human tumors and potential roles in carcinogenesis. Mutat. Res. 533, 201–209 (2003).

56. Jin, R. et al. Metallothionein 2A expression is associated with cell proliferation in breast cancer. Carcinogenesis 23, 81–86 (2002).

57. Pereira, H. et al. Metallothionein expression in human breast cancer. The Breast vol. 1 159–160 (1992).

58. Pedersen, M. Ø., Larsen, A., Stoltenberg, M. & Penkowa, M. The role of metallothionein in oncogenesis and cancer prognosis. Prog. Histochem. Cytochem. 44, 29–64 (2009).

59. Laughney, A. M. et al. Regenerative lineages and immune-mediated pruning in lung cancer metastasis. Nat. Med. 26, 259–269 (2020).

60. Marjanovic, N. D. et al. Emergence of a High-Plasticity Cell State during Lung Cancer Evolution. Cancer Cell 38, 229–246.e13 (2020).

61. Maynard, A. et al. Therapy-Induced Evolution of Human Lung Cancer Revealed by Single-Cell RNA Sequencing. Cell 182, 1232–1251.e22 (2020).

62. Hao, D. et al. Integrated Analysis Reveals Tubal-and Ovarian-Originated Serous Ovarian Cancer and Predicts Differential Therapeutic Responses. Clin. Cancer Res. 23, 7400–7411 (2017).

63. Zhang, S. et al. Both fallopian tube and ovarian surface epithelium are cells-of-origin for high-grade serous ovarian carcinoma. Nat. Commun. 10, 5367 (2019).

64. Aiello, N. M. et al. EMT Subtype Influences Epithelial Plasticity and Mode of Cell Migration. Dev. Cell 45, 681–695.e4 (2018).

65. Fischer, K. R. et al. Epithelial-to-mesenchymal transition is not required for lung metastasis but contributes to chemoresistance. Nature vol. 527 472–476 (2015).

66. Hoadley, K. A. et al. Cell-of-Origin Patterns Dominate the Molecular Classification of 10,000 Tumors from 33 Types of Cancer. Cell 173, 291–304.e6 (2018).

67. Kalluri, R. & Weinberg, R. A. The basics of epithelial-mesenchymal transition. J. Clin. Invest. 119, 1420–1428 (2009).

68. Cook, D. P. & Vanderhyden, B. C. Transcriptional census of epithelial-mesenchymal plasticity in cancer. bioRxiv 2021.03.05.434142 (2021) doi:10.1101/2021.03.05.434142.

69. Aibar, S. et al. SCENIC: single-cell regulatory network inference and clustering. Nat. Methods 14, 1083–1086 (2017).

70. Pu, W. et al. Single-cell transcriptomic analysis of the tumor ecosystems underlying initiation and progression of papillary thyroid carcinoma. Nat. Commun. 12, 6058 (2021).

71. Hayashi, A. et al. A unifying paradigm for transcriptional heterogeneity and squamous features in pancreatic ductal adenocarcinoma. Nature Cancer vol. 1 59–74 (2020).

72. Collisson, E. A. et al. Subtypes of pancreatic ductal adenocarcinoma and their differing responses to therapy. Nat. Med. 17, 500–503 (2011).

73. Moffitt, R. A. et al. Virtual microdissection identifies distinct tumor-and stroma-specific subtypes of pancreatic ductal adenocarcinoma. Nat. Genet. 47, 1168–1178 (2015).

74. Baylor, S. M. & Berg, J. W. Cross-classification and survival characteristics of 5,000 cases of cancer of the pancreas. J. Surg. Oncol. 5, 335–358 (1973).

75. Al-Shehri, A., Silverman, S. & King, K. M. Squamous Cell Carcinoma of the Pancreas. Current Oncology vol. 15 293–297 (2008).

76. Hanahan, D. & Weinberg, R. A. The Hallmarks of Cancer. Cell vol. 100 57–70 (2000).

77. Hanahan, D. & Weinberg, R. A. Hallmarks of Cancer: The Next Generation. Cell vol. 144 646–674 (2011).

78. Archetti, M. & Pienta, K. J. Cooperation among cancer cells: applying game theory to cancer. Nat. Rev. Cancer 19, 110–117 (2019).

79. Diamond, M. S. et al. Type I interferon is selectively required by dendritic cells for immune rejection of tumors. J. Exp. Med. 208, 1989–2003 (2011).

80. Deng, L. et al. STING-Dependent Cytosolic DNA Sensing Promotes Radiation-Induced Type I Interferon-Dependent Antitumor Immunity in Immunogenic Tumors. Immunity 41, 843–852 (2014).

81. Ng, K. W., Marshall, E. A., Bell, J. C. & Lam, W. L. cGAS-STING and Cancer: Dichotomous Roles in Tumor Immunity and Development. Trends Immunol. 39, 44–54 (2018).

82. Williams, J. B. et al. Tumor heterogeneity and clonal cooperation influence the immune selection of IFN-γ-signaling mutant cancer cells. Nat. Commun. 11, 602 (2020).

83. Kinker, G. S. et al. Pan-cancer single cell RNA-seq uncovers recurring programs of cellular heterogeneity. doi:10.1101/807552.

84. Ståhl, P. L. et al. Visualization and analysis of gene expression in tissue sections by spatial transcriptomics. Science 353, 78–82 (2016).

85. Francis, K. & Palsson, B. O. Effective intercellular communication distances are determined by the relative time constants for cyto/chemokine secretion and diffusion. Proc. Natl. Acad. Sci. U. S. A. 94, 12258–12262 (1997).

86. Stuart, T. et al. Comprehensive Integration of Single-Cell Data. Cell 177, 1888–1902.e21 (2019).

87. Weis, S. M. & Cheresh, D. A. Tumor angiogenesis: molecular pathways and therapeutic targets. Nature Medicine vol. 17 1359–1370 (2011).

88. Viallard, C. & Larrivée, B. Tumor angiogenesis and vascular normalization: alternative therapeutic targets. Angiogenesis 20, 409–426 (2017).

89. Solinas, G., Germano, G., Mantovani, A. & Allavena, P. Tumor-associated macrophages (TAM) as major players of the cancer-related inflammation. J. Leukoc. Biol. 86, 1065–1073 (2009).

90. Zhang, M. et al. A high M1/M2 ratio of tumor-associated macrophages is associated with extended survival in ovarian cancer patients. J. Ovarian Res. 7, 19 (2014).

91. Yuan, A. et al. Opposite Effects of M1 and M2 Macrophage Subtypes on Lung Cancer Progression. Sci. Rep. 5, 14273 (2015).

92. Pinto, M. L. et al. The Two Faces of Tumor-Associated Macrophages and Their Clinical Significance in Colorectal Cancer. Front. Immunol. 10, 1875 (2019).

93. Cheng, S. et al. A pan-cancer single-cell transcriptional atlas of tumor infiltrating myeloid cells. Cell 184, 792–809.e23 (2021).

94. Hara, T. et al. Interactions between cancer cells and immune cells drive transitions to mesenchymal-like states in glioblastoma. Cancer Cell 39, 779–792.e11 (2021).

95. Ohkuri, T. et al. Intratumoral administration of cGAMP transiently accumulates potent macrophages for anti-tumor immunity at a mouse tumor site. Cancer Immunology, Immunotherapy vol. 66 705–716 (2017).

96. Black, S. et al. CODEX multiplexed tissue imaging with DNA-conjugated antibodies. Nat. Protoc. 16, 3802–3835 (2021).

97. Rozenblatt-Rosen, O. et al. The Human Tumor Atlas Network: Charting Tumor Transitions across Space and Time at Single-Cell Resolution. Cell 181, 236–249 (2020).

98. Weinberg, R. A. The Biology of Cancer. (Garland Pub, 2007).

99. Sottoriva, A. et al. A Big Bang model of human colorectal tumor growth. Nat. Genet. 47, 209–216 (2015).

100. Zaidi, M. R. & Merlino, G. The two faces of interferon-γ in cancer. Clin. Cancer Res. 17, 6118–6124 (2011).

101. Fisher, R., Pusztai, L. & Swanton, C. Cancer heterogeneity: implications for targeted therapeutics. British Journal of Cancer vol. 108 479–485 (2013).

102. Klein, A. M. et al. Droplet barcoding for single-cell transcriptomics applied to embryonic stem cells. Cell 161, 1187–1201 (2015).

103. Baron, M. et al. A Single-Cell Transcriptomic Map of the Human and Mouse Pancreas Reveals Inter-and Intra-cell Population Structure. Cell Syst 3, 346–360.e4 (2016).

104. Hafemeister, C. & Satija, R. Normalization and variance stabilization of single-cell RNA-seq data using regularized negative binomial regression. Genome Biol. 20, 296 (2019).

105. Mabbott, N. A., Baillie, J. K., Brown, H., Freeman, T. C. & Hume, D. A. An expression atlas of human primary cells: inference of gene function from coexpression networks. BMC Genomics 14, 632 (2013).

106. Galgano, M. T., Hampton, G. M. & Frierson, H. F. Comprehensive analysis of HE4 expression in normal and malignant human tissues. Modern Pathology vol. 19 847–853 (2006).

107. Chen, L., O’Bryan, J. P., Smith, H. S. & Liu, E. Overexpression of matrix Gla protein mRNA in malignant human breast cells: isolation by differential cDNA hybridization. Oncogene 5, 1391–1395 (1990).

108. Kosanam, H. et al. Laminin, gamma 2 (LAMC2): a promising new putative pancreatic cancer biomarker identified by proteomic analysis of pancreatic adenocarcinoma tissues. Mol. Cell. Proteomics 12, 2820–2832 (2013).

109. Zheng, B. et al. TM4SF1 as a prognostic marker of pancreatic ductal adenocarcinoma is involved in migration and invasion of cancer cells. Int. J. Oncol. 47, 490–498 (2015).

110. Jothy, S., Yuan, S. Y. & Shirota, K. Transcription of carcinoembryonic antigen in normal colon and colon carcinoma. In situ hybridization study and implication for a new in vivo functional model. Am. J. Pathol. 143, 250–257 (1993).

111. Jing, X., Piao, Y.-F., Liu, Y. & Gao, P.-J. Beta2-GPI: a novel factor in the development of hepatocellular carcinoma. J. Cancer Res. Clin. Oncol. 136, 1671–1680 (2010).

112. Landers, K. A. et al. Identification of claudin-4 as a marker highly overexpressed in both primary and metastatic prostate cancer. Br. J. Cancer 99, 491–501 (2008).

113. Liao, S. Y., Aurelio, O. N., Jan, K., Zavada, J. & Stanbridge, E. J. Identification of the MN/CA9 protein as a reliable diagnostic biomarker of clear cell carcinoma of the kidney. Cancer Res. 57, 2827–2831 (1997).

114. Allander, S. V. et al. Gastrointestinal stromal tumors with KIT mutations exhibit a remarkably homogeneous gene expression profile. Cancer Res. 61, 8624–8628 (2001).

115. West, R. B. et al. The Novel Marker, DOG1, Is Expressed Ubiquitously in Gastrointestinal Stromal Tumors Irrespective of KIT or PDGFRA Mutation Status. The American Journal of Pathology vol. 165 107–113 (2004).

116. Gaujoux, R. & Seoighe, C. A flexible R package for nonnegative matrix factorization. BMC Bioinformatics 11, 367 (2010).

117. Carmona-Saez, P., Pascual-Marqui, R. D., Tirado, F., Carazo, J. M. & Pascual-Montano, A. Biclustering of gene expression data by Non-smooth Non-negative Matrix Factorization. BMC Bioinformatics 7, 78 (2006).

118. igraph – Network analysis software. http://igraph.org.

119. Subramanian, A. et al. Gene set enrichment analysis: a knowledge-based approach for interpreting genome-wide expression profiles. Proc. Natl. Acad. Sci. U. S. A. 102, 15545–15550 (2005).

120. Franzén, O., Gan, L.-M. & Björkegren, J. L. M. PanglaoDB: a web server for exploration of mouse and human single-cell RNA sequencing data. Database 2019, (2019).

121. Ibrahim, M. M. & Kramann, R. genesorteR: Feature Ranking in Clustered Single Cell Data. bioRxiv 676379 (2019) doi:10.1101/676379.

122. Thorsson, V. et al. The Immune Landscape of Cancer. Immunity 48, 812–830.e14 (2018).

123. CRAN - Package nnls. https://CRAN.R-project.org/package=nnls.

124. Elyada, E. et al. Cross-Species Single-Cell Analysis of Pancreatic Ductal Adenocarcinoma Reveals Antigen-Presenting Cancer-Associated Fibroblasts. Cancer Discov. 9, 1102–1123 (2019).

125. Durinck, S. et al. BioMart and Bioconductor: a powerful link between biological databases and microarray data analysis. Bioinformatics 21, 3439–3440 (2005).

